# Raw signal segmentation for estimating RNA modification from Nanopore direct RNA sequencing data

**DOI:** 10.1101/2024.01.11.575207

**Authors:** Guangzhao Cheng, Aki Vehtari, Lu Cheng

## Abstract

Estimating RNA modifications from Nanopore direct RNA sequencing data is a critical task for the RNA research community. However, current computational methods often fail to deliver satisfactory results due to inaccurate segmentation of the raw signal. We have developed a new method, SegPore, which leverages a molecular jiggling translocation hypothesis to improve raw signal segmentation. SegPore is a pure white-box model with enhanced interpretability, significantly reducing structured noise in the raw signal. We demonstrate that SegPore outperforms state-of-the-art methods, such as Nanopolish and Tombo, in raw signal segmentation across three large benchmark datasets. Moreover, the improved signal segmentation achieved by SegPore enables SegPore+m6Anet to deliver state-of-the-art performance in site-level m6A identification. Additionally, SegPore surpasses baseline methods like CHEUI in single-molecule level m6A identification.

## INTRODUCTION

RNA modifications play important roles in different diseases, such as Acute Myeloid Leukemia (1) and Fragile X Syndrome (2), as well as fundamental biological processes like cell differentiation (3,4) and immune response (5). To date, researchers have identified over 150 different types of RNA modifications (6-9), highlighting the complexity and diversity of RNA regulation. These modifications are essential for proper RNA function, including maintaining secondary structures and facilitating accurate protein synthesis, such as the role of tRNA modifications in decoding mRNA codons (10).

The practical applications of RNA modifications are far-reaching. For instance, N1-methylpseudouridine (m1Ψ) has been used to enhance the efficacy of COVID-19 mRNA vaccines, highlighting their therapeutic potential (11). However, identifying and characterizing RNA modifications presents significant challenges due to the limitations of existing experimental techniques.

Traditional methods for RNA modification detection, such as MeRIP-Seq (12), miCLIP (13), and m6ACE-Seq (14), rely on immunoprecipitation techniques that use modification-specific antibodies. While effective, these methods have several drawbacks. First, each method requires a separate antibody for each RNA modification, making it difficult to study multiple modifications simultaneously. Additionally, these techniques can only infer modification locations from next-generation sequencing (NGS) data, rather than measuring the modifications directly. As a result, they struggle to provide single-molecule resolution, limiting their accuracy and scope.

Recent advancements in direct RNA sequencing (DRS) by Oxford Nanopore Technologies (ONT) offer a promising alternative. DRS allows for the direct measurement of electrical currents as RNA molecules translocate through a nanopore, providing a potential avenue for detecting RNA modifications at the single-molecule level. Two versions of the direct RNA sequencing (DRS) kits are available: RNA002 and RNA004. Unless otherwise specified, this study focuses on RNA002 data. In this technique, five nucleotides (5-mers) reside in the nanopore at a time, and each 5-mer generates a characteristic current signal based on its unique sequence and chemical properties (16). By analyzing these signals, it is possible to infer the original RNA sequence and detect modifications like N6-methyladenosine (m6A).

The general workflow of Nanopore direct RNA sequencing (DRS) data analysis is as follows. First, the raw electrical signal from a read is basecalled using tools such as Guppy or Dorado (https://github.com/nanoporetech), which produce the nucleotide sequence of the RNA molecule. However, these basecalled sequences do not include the precise start and end positions of each ribonucleotide (or k-mer) in the signal. Because basecalling errors are common, the sequences are typically mapped to a reference genome or transcriptome using minimap2 (15) to recover the correct reference sequence. Next, tools such as Nanopolish (16,17) and Tombo (18) align the raw signal to the reference sequence to determine which portion of the signal corresponds to each k-mer. We define this process as the segmentation and alignment task (abbreviated as the segmentation task), which is referred to as “eventalign” in Nanopolish. Based on this alignment, Nanopolish extracts various features—such as the start and end positions, mean, and standard deviation of the signal segment corresponding to a k-mer. This signal segment or its derived features is referred to as an **“**event**”** in Nanopolish. The resulting events serve as input for downstream RNA modification detection tools such as m6Anet (19) and CHEUI (20).

However, significant computational challenges remain. Segmenting the raw current signal into distinct 5-mers and distinguishing between normal nucleotides and their modified counterparts is a complex task. Current methods, such as Nanopolish, employ change-point detection methods to segment the signal and use dynamic programming methods and Hidden Markov Models (HMM) to align the derived segments to the reference sequence, but they are prone to noise and inaccuracies, which degrade performance in downstream tasks like RNA modification prediction. The root of this issue lies in the fact that these methods do not accurately model the physical process of Nanopore sequencing, particularly the dynamics of the motor protein that drives RNA through the pore. As a result, the segmentation process is not well-aligned with the actual translocation mechanics, leading to signal noise that hinders precise modification detection.

SegPore is a novel tool for direct RNA sequencing (DRS) signal segmentation and alignment, designed to overcome key limitations of existing approaches. By explicitly modeling motor protein dynamics during RNA translocation with a Hierarchical Hidden Markov Model (HHMM), SegPore segments the raw signal into small, biologically meaningful fragments, each corresponding to a k-mer sub-state, which substantially reduces noise and improves segmentation accuracy. After segmentation, these fragments are aligned to the reference sequence and concatenated into larger events, analogous to Nanopolish’s “eventalign” output, which serve as the foundation for downstream analyses. Moreover, the “eventalign” results produced by SegPore enhance interpretability in RNA modification estimation. While deep learning–based tools such as m6Anet classify RNA modifications using complex, non-transparent features (see Supplementary Fig. S5), SegPore employs a simple Gaussian Mixture Model (GMM) to distinguish modified from unmodified nucleotides based on baseline current levels. This transparent modeling approach improves confidence in the predictions and makes SegPore particularly well-suited for biological applications where interpretability is essential.

By introducing SegPore, we bridge the gaps left by traditional tools. SegPore utilizes a hierarchical hidden Markov model (HHMM) for more precise segmentation and combines it with signal alignment and a GMM for RNA modification prediction, offering both greater accuracy and interpretability. This integrated approach enables robust m6A modification detection at the single-molecule level, facilitating reliable, transparent predictions for both site-specific and molecule-wide m6A modifications.

## MATERIALS AND METHODS

### SegPore workflow

#### Workflow overview

An overview of SegPore is illustrated in Figure 1A, which outlines its five-step process. The output of Step 3 is the “events”, which is analogous to the output generated by the Nanopolish (v0.14.0) “eventalign” command and can be used as input for downstream models such as m6Anet. An “event” refers to a segment of the raw signal that is aligned to a specific k-mer on a read, along with its associated features such as start and end positions, mean current, standard deviation, and other relevant statistics. Step 4 allows for direct modification prediction at both the site and single-molecule levels. Notably, a key feature of SegPore is the k-mer parameter table, which defines the mean and standard deviation for each k-mer in either an unmodified or modified state. During the training phase, Steps 3~5 are iterated multiple times to stabilize the parameters in the k-mer table, which are subsequently fixed and applied for modification prediction on the test data. A detailed description of the model is provided in Supplementary Note 1. Unless otherwise noted, the following analysis focuses on RNA002 data, using 5-mers rather than k-mers.

**Figure 1.**
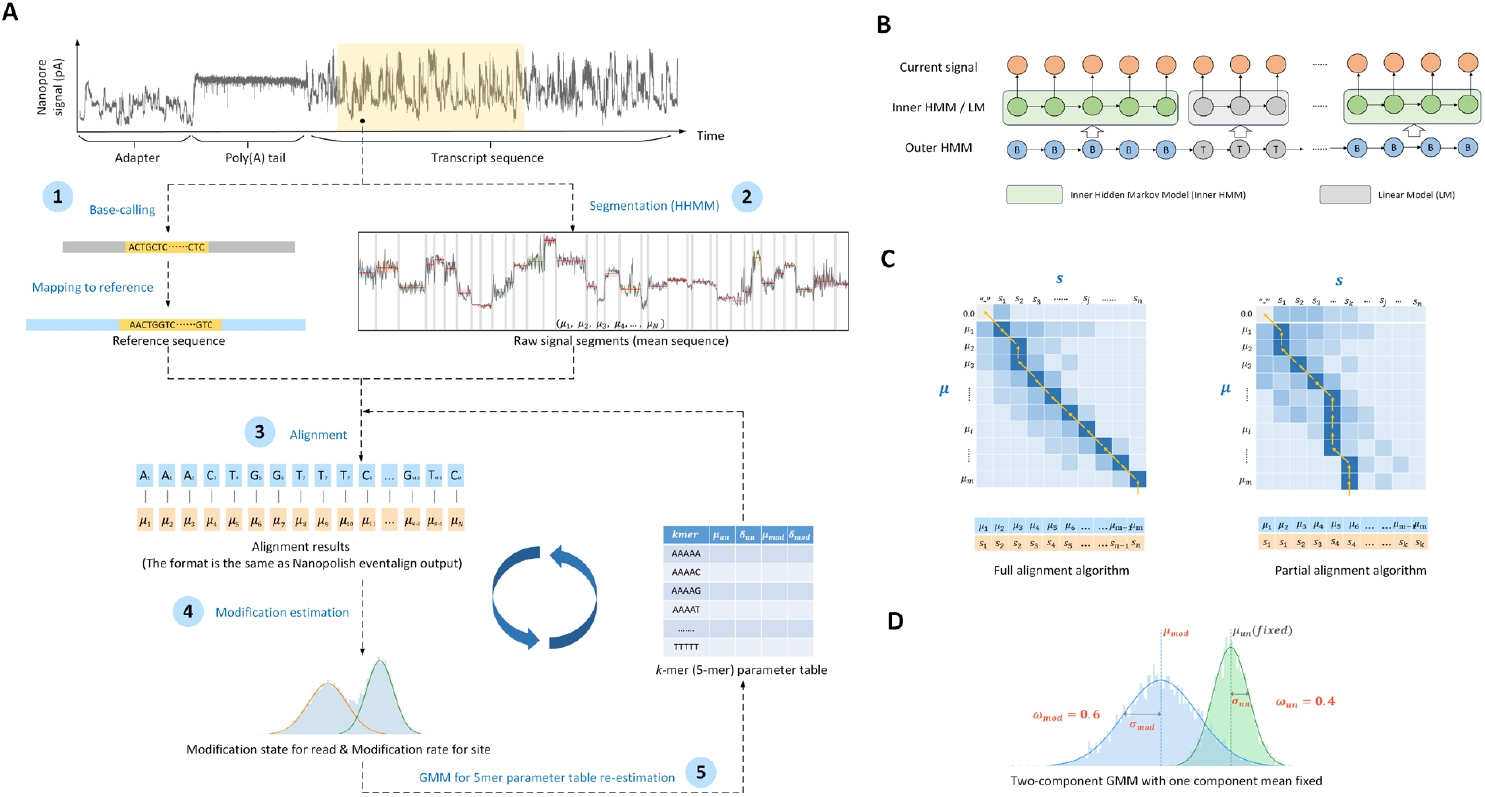
SegPore workflow. (A) General workflow. The workflow consists of five steps: (1) First, raw current signals are basecalled and mapped using Guppy and Minimap2. The raw current signal fragments are paired with the corresponding reference RNA sequence fragments using Nanopolish. (2) Next, the raw current signal of each read is segmented using a hierarchical hidden Markov model (HHMM), which provides an estimated mean (*μ*_*i*_) for each segment. (3) These segments are then aligned with the 5-mer list of the reference sequence fragment using a full/partial alignment algorithm, based on a 5-mer parameter table. For example, *A*_*j*_ denotes the base “*A*” at the *j*-th position on the reference. In this example, *A*_1_ and *A*_2_ refer to the first and second occurrences of “*A*” in the reference sequence, respectively. Accordingly, *μ*_1_ and *μ*_2_ are aligned to *A*_1_, while *μ*_3_ is aligned to *A*_2_. (4) All signals aligned to the same 5-mer across different genomic locations are pooled together, and a two-component Gaussian Mixture Model (GMM) is used to predict the modification at the site-level or single-molecule level. One component of the GMM represents the unmodified state, while the other represents the modified state. (5) GMM is used to re-estimate the 5-mer parameter table. (B) Hierarchical hidden Markov model (HHMM). The outer HMM segments the current signal into alternating base and transition blocks. The inner HMM approximates the emission probability of a base block by considering neighboring 5-mers. A linear model is used to approximate the emission probability of a transition block. (C) Full/partial alignment algorithms. Rows represent the estimated means of base blocks from the HHMM, and columns represent the 5-mers of the reference sequence. Each 5-mer can be aligned with multiple estimated means from the current signal. (D) Gaussian mixture model (GMM) for estimating modification states. The GMM consists of two components: the green component models the unmodified state of a 5-mer, and the blue component models the modified state. Each component is described by three parameters: mean (*μ*), standard deviation (*σ*), and weight (*ω*).

#### Preprocessing

We begin by performing basecalling on the input fast5 file using Guppy (v6.0.1), which converts the raw signal data into ribonucleotide sequences. Next, we align the basecalled sequences to the reference genome using Minimap2 (v2.28-r1209) (15), generating a mapping between the reads and the reference sequences. Nanopolish provides two independent commands: “polya” and “eventalign”. The “polya” command identifies the adapter, poly(A) tail, and transcript region in the raw signal, which we refer to as the poly(A) detection results. The raw signal segment corresponding to the poly(A) tail is used to standardize the raw signal for each read. The “eventalign” command aligns the raw signal to a reference sequence, assigning a signal segment to individual k-mers in the reference. It also computes summary statistics (e.g., mean, standard deviation) from the signal segment for each k-mer. Each k-mer together with its corresponding signal features is termed an event. These event features are then passed into downstream tools such as m6Anet and CHEUI for RNA modification detection. For full transcriptome analysis (Fig. 3), we extract the aligned raw signal segment and reference sequence segment from Nanopolish’s events for each read by using the first and last events as start and end points. For *in vitro* transcription (IVT) data with a known reference sequence (Fig. 4), we extract the raw signal segment corresponding to the transcript region for each input read based on Nanopolish’s poly(A) detection results.

Due to inherent variability between nanopores in the sequencing device, the baseline levels and standard deviations of k-mer signals can differ across reads, even for the same transcript. To standardize the signal for downstream analyses, we extract the raw current signal segments corresponding to the poly(A) tail of each read. Since the poly(A) tail provides a stable reference, we standardize the raw current signals for each read, ensuring that the mean and standard deviation are consistent across the poly(A) tail region. This step is crucial for reducing variability between different reads and ensuring more accurate signal segmentation and modification prediction in subsequent steps. See Section 3 of Supplementary Note 1 for more details.

#### Signal segmentation via hierarchical Hidden Markov model

The RNA translocation hypothesis (see details in the first section of *Results*) naturally leads to the use of a hierarchical Hidden Markov Model (HHMM) to segment the raw current signal. As shown in Figure 1B, the HHMM consists of two layers. The outer HMM divides the raw signal into alternating base and transition blocks, represented by hidden states “B” (base) and “T” (transition). Within each base block, the inner HMM models the current signal at a more granular level.

The inner HMM includes four hidden states: “prev,” “next,” “curr,” and “noise.” These correspond to the previous, next, and current 5-mer in the pore, while “noise” refers to random fluctuations. Each raw current measurement is emitted from one of these hidden states, providing a detailed model of the signal within the base blocks. A linear model with a large absolute slope is used to represent sharp changes in the transition blocks.

To segment the signal, we first model the likelihood of the HHMM. Given the raw current signal ***y*** of a read, the hidden states of the outer hidden HMM are denoted by ***g. y*** and ***g*** are divided into 2*K* + 1 blocks ***c***, where ***y***^(*k*)^, ***g***^(*k*)^ correspond to *k* th block and *c* = (*c*_1_, *c*_2_, …, *c*_*k*_, …, *c*_2*K*+1_), *c*_*k*_ ∈ {*B, T*}. Blocks with odd indices (*k* = 1, 3, 5, …, 2*K* + 1) are base blocks, while those with even indices (*k* = 2, 4, …, 2*K*) are transition blocks. The likelihood of the HHMM is given by

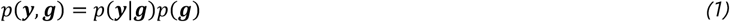

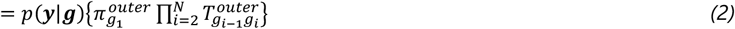

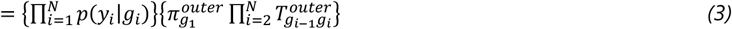

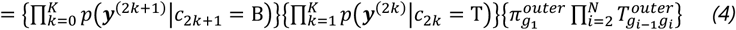

where 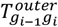 is the transition matrix of the outer HMM and 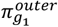 is the probability for the first hidden state. The first part of the Eq. 2 are emission probabilities, and the second part are the transition probabilities. It is not possible to directly compute the emission probabilities of the outer HMM (Eq. 3) since there exist dependencies for the current signal measurements within a base or transition block. Therefore, we use the inner HMM and linear model (Eq. 4) to handle the dependencies and approximate emission probabilities.

The inner HMM models transitions between the “prev,” “next,” “curr,” and “noise” states. For the “prev,” “next,” and “curr” states, a Gaussian distribution is used to model the emission probabilities, while a uniform distribution models the “noise” state. The forward-backward algorithm is used to compute the marginal likelihood of the inner HMM, which approximates the emission probabilities for base blocks. For transition blocks, a standard linear model computes the emission probabilities.

For any given ***g***, we can calculate the joint likelihood (Eq. 1). We enumerate different configurations and select the one with the highest likelihood. This process segments the raw current signal into alternating base and transition blocks, where one or more base blocks may correspond to a single 5-mer. Each base block is characterized by its mean and standard deviation, which are used as input for downstream alignment tasks.

Due to the large size of fast5 files (which can reach terabytes), parameter inference for this model is computationally intensive. To address this, we developed a GPU-based inference algorithm that significantly accelerates the process. More details on this algorithm can be found in Supplementary Note 2.

In summary, the HHMM allows us to accurately segment the raw current signal, providing estimates of the mean and variance for each base block, which are crucial for downstream analyses such as RNA modification prediction.

#### 5-mer & 9-mer parameter table

We downloaded the k-mer models “r9.4_180mv_70bps_5-mer_RNA” from the ONT GitHub repository (https://github.com/nanoporetech/kmer_models) for RNA002 data. The columns labeled “level_mean” and “level_stdv” in these models were used as the mean and standard deviation values for the unmodified 5-mers. These values serve as the initial parameters in the 5-mer parameter table for SegPore, which we refer to as the *ONT 5-mer parameter table*. In the RNA004 data analysis, we obtained the 9-mer parameter table from the source code of f5c (version 1.5). Specifically, we used the array named “rna004_130bps_u_to_t_rna_9mer_template_model_builtin_data” from the following file: https://raw.githubusercontent.com/hasindu2008/f5c/refs/heads/master/src/model.h (accessed on 17 October 2025).

The initialization of the k-mer parameter table is a critical step in SegPore’s workflow. By leveraging ONT’s established k-mer models, we ensure that the initial estimates for unmodified k-mers are grounded in empirical data. These initial estimates are refined through SegPore’s iterative parameter estimation process, enabling the model to accurately differentiate between modified and unmodified k-mers during segmentation and modification prediction tasks. The refined k-mer parameters also provide a foundation for downstream analysis, such as alignment and m6A identification.

#### Alignment of raw signal segment with reference sequence

After segmenting the raw current signal of a read into base and transition blocks using HHMM, we align the means of base blocks with the 5-mer list of the reference sequence.

The alignment process involves three main cases:

1. Base block matching with a 5-mer: In this case, a base block aligns with a 5-mer, which is modeled by a Gaussian distribution. The 5-mer may exist in either an unmodified or modified state, and the corresponding Gaussian parameters are retrieved from the 5-mer parameter table.
2. Base block matching with an insertion: Here, the base block aligns with an indel (“-”), indicating an inserted nucleotide in the read.
3. Deletion in the read: In this case, an indel (0.0) matches with a 5-mer, representing a deleted nucleotide in the read.

The alignment score function models these matching cases as follows. For the first case (a base block matching a 5-mer), we calculate the probability that the base block mean is sampled from either the unmodified or modified 5-mer’s Gaussian distribution. The larger of the two probabilities is used as the match score. For the second and third cases, the alignment is treated as noise, and a fixed uniform distribution is used to calculate the match score.

An important distinction from classical global alignment algorithms (Needleman–Wunsch algorithm) is that one or multiple base blocks may align with a single 5-mer. Given the base block means ***μ*** = (*μ*_1_, *μ*_2_, …, *μ*_*i*_, …, *μ*_*m*_) and 5-mer list *s* = (*s*_1_, *s*_2_, …, *s*_*j*_, …, *s*_*n*_), we define (*m* + 1) × (*n* + 1) the score matrix as ***M*** (Fig. 1C). The first row and column of the matrix are reserved for indels (“0.0” and “-”), representing insertions or deletions in the base blocks or 5-mers, respectively.

We denote the score function by *f*. The recursion formula of the dynamic programming alignment algorithm is given by

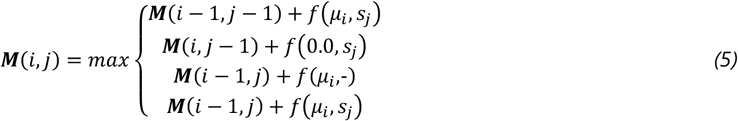

, where *f* is the score function. It can be seen that we can still align *μ*_*i*_ with *s*_*j*_ after we have aligned *μ*_*i*−1_ with *s*_*j*_, which fulfills the special consideration that one or multiple base blocks might be aligned with one 5-mer.

We implement two types of alignment algorithms (Fig. 1C) based on the score matrix ***M***:

1. Full Alignment Algorithm: This algorithm aligns the full list of base block means with the full list of 5-mers, similar to classical global alignment. It traces back from the (*m* + 1, *n* + 1) position in the score matrix.
2. Partial Alignment Algorithm: This aligns the full list of base blocks with the initial part of the 5-mer list, with no indels allowed in the base block means or the 5-mer list. The trace back starts from the maximum value in the last row of the score matrix.

A detailed description of both alignment algorithms is provided in Supplementary Note 1. The output of the alignment algorithm is an alignment that pairs the base blocks with the 5-mers from the reference sequence for each read (Fig. 1C). Base blocks aligned to the same 5-mer are concatenated into a single raw signal segment (referred to as an “event”), from which various features—such as start and end positions, mean current, and standard deviation—are extracted. Detailed derivation of the mean and standard deviation is provided in Section 5.3 in Supplementary Note 1. In the remainder of this paper, we refer to these resulting events as the output of eventalign analysis, which also represents the final output of the segmentation and alignment task.

#### Modification prediction

After obtaining the eventalign results, we estimate the modification state of each motif using the 5-mer parameter table. Specifically, for each 5-mer, we compare the probability that its mean is sampled from either the modified or unmodified 5-mer’s Gaussian distribution. If the probability under the modified 5-mer distribution is higher, the 5-mer is predicted to be in the modification state, and vice versa for the unmodified state.

To estimate the overall modification rate at a specific genomic location, we pool all reads that map to that location on the reference sequence. The modification rate is calculated as the proportion of reads that are predicted to be in the modification state at that location. A detailed description of the modification state prediction process can be found in Supplementary Note 1.

#### GMM for 5-mer parameter table re-estimation

To improve alignment accuracy and enhance modification predictions, we use a Gaussian Mixture Model (GMM) to iteratively fine-tune the 5-mer parameter table. As illustrated in Figure 1A, the rows of the 5-mer parameter table represent the 5-mers, while the columns provide the mean and standard deviation for both the unmodified and modified states. We denote a 5-mer by *s*, with its relevant parameters as *μ*_*s,un*_, *δ*_*s,un*_, *μ*_*s,mod*_, *δ*_*s,mod*_.

Using the alignment results from all reads, we collect the base block means aligned to the same 5-mer (denoted by *s*) across different reads and genomic locations, particularly those with high modification rates. A two-component GMM is then fit to these base blocks. In this process, the mean of the first component is fixed to *μ*_*s,un*_. From the GMM, we update the parameters *δ*_*s,un*_, *ω*_*s,un*_, *μ*_*s,mod*_, *δ*_*s,mod*_, and *ω*_*s,mod*_, where *ω*_*s,un*_, *ω*_*s,mod*_ represent the weights for the unmodified and modified components, respectively. Afterward, we manually adjust the 5-mer parameter table using heuristics to ensure that the modified 5-mer distribution is significantly distinct from the unmodified distribution (see details in Section 7 of Supplementary Note 1).

This re-estimation process is only performed on the training data. The initial 5-mer parameter table is based on the table provided by ONT. After each iteration of updating the table, we rerun the SegPore workflow from the alignment step onward. Typically, the process is repeated three to five times until the 5-mer parameter table stabilizes (when the average change of mean values of all 5-mers is less than 5e-3). Once a stabilized 5-mer parameter table is estimated from the training data, it is used for RNA modification estimation in the test data without further updates. A more detailed description of the GMM re-estimation process is provided in Section 6 of Supplementary Note 1.

### m6A site level benchmark

The HEK293T wild-type (WT) samples were downloaded from the ENA database (accession number PRJEB40872), while the HCT116 samples were obtained from ENA under accession PRJEB44348. The reference sequence (Homo_sapiens.GRCh38.cdna.ncrna_wtChrIs_modified.fa) was downloaded from https://doi.org/10.5281/zenodo.4587661. Ground truth data were sourced from Supplementary Data 1 of Pratanwanich, P.N. et al. (21). Fast5 files for the test dataset (mESC WT samples, mESCs_Mettl3_WT_fast5.tar.gz) (22) were retrieved from the NCBI Sequence Read Archive (SRA) under accession SRP166020.

During training, we initialized the 5-mer parameter table using ONT’s data. The standard SegPore workflow was executed on the training data (HEK293T WT samples), using the full alignment algorithm. The 5-mer parameter table estimation was iterated five times. For mapping, reads were first aligned to the cDNA and subsequently converted to genomic locations using Ensembl GTF file (GRCh38, v9). The same 5-mer across different genomic locations was pooled together. A 5-mer was considered significantly modified if its read coverage exceeded 1,500 and the distance between the means of the two Gaussian components in the GMM was greater than 5 picoamperes (pA). As a result, modification parameters were specified for ten significant 5-mers, as illustrated in Supplementary Fig. S2A.

With the estimated 5-mer parameter table from the training data, we then ran the SegPore workflow on the test data. The transcript sequences from GENCODE release version M18 were used as the reference sequence for mapping, with the corresponding GTF file (gencode.vM18.chr_patch_hapl_scaff.annotation.gtf) downloaded from GENCODE to convert transcript locations into genomic coordinates. It is important to note that the 5-mer parameter table was not re-estimated for the test data. Instead, modification states for each read were directly estimated using the fixed 5-mer parameter table. Due to the differences between human (Supplementary Fig. S2A) and mouse (Supplementary Fig. S2B), only six 5-mers were found to have m6A annotations in the test data’s ground truth (Supplementary Fig. S2C). For a genomic location to be identified as a true m6A modification site, it had to correspond to one of these six common 5-mers and have a read coverage of greater than 20. SegPore derived the ROC and PR curves for benchmarking based on the modification rate at each genomic location.

In the SegPore+m6Anet analysis, we fine-tuned the m6Anet model using SegPore’s eventalign results to demonstrate improved m6A identification. We started with the pre-trained m6Anet model (available at https://github.com/GoekeLab/m6anet, model version: HCT116_RNA002) and fine-tuned it using the eventalign results from HCT116 samples. SegPore’s eventalign output provided the pairing between each genomic location and its corresponding raw signal segment, which allowed us to extract normalized features such as the normalized mean *μ*_*i*_, standard deviation *σ*_*i*_, dwell time *l*_*i*_ (number of data points in the event). For genomic location *i*, m6Anet extracts a feature vector *x*_*i*_ = {*μ*_*i*−1_, *σ*_*i*−1_, *l*_*i*−1_, *μ*_*i*_, *σ*_*i*_, *l*_*i*_, *μ*_*i*+1_, *σ*_*i*+1_, *l*_*i*+1_}, which was used as input for m6Anet. Feature vectors from a randomly selected 80% of the genomic locations were used for training, while the remaining 20% were set aside for validation. We ran 100 epochs during fine-tuning, selecting the model that performed best on the validation set.

Ground truth data and the performance of other methods (Tombo v1.5.1, Nanom6A v2.0, m6Anet v1.0, and Epinano v1.2.0) on the mESC dataset were provided through personal communications with Prof. Luo Guanzheng, the corresponding author of the benchmark study referenced (23).

### m6A single molecule level benchmark

The benchmark IVT data for single molecule m6A identification was downloaded from NCBI-SRA with accession number SRP166020. CHEUI was used for the benchmark. For the ground truth, every A in every read of the IVT_m6A sample is treated as a m6A modification and every A in every read of the IVT_normalA is treated as a normal A. Detailed methods for calculating modification probability at single molecule level were provided in Supplementary Note 1, Section 6.1-6.2.

## RESULTS

### RNA translocation hypothesis

Accurate segmentation of raw current signals in direct RNA sequencing remains a major challenge, largely because the precise dynamics of RNA translocation through the pore are not fully understood. To address this, we propose a hypothesis that better reflects the physical movement of the RNA molecule. In traditional basecalling algorithms such as Guppy and Albacore, we implicitly assume that the RNA molecule is translocated through the pore by the motor protein in a monotonic fashion, i.e., the RNA is pulled through the pore unidirectionally. In the DNN training process of Guppy and Albacore, we try to align the current signal with the reference RNA sequence. The alignment is unidirectional, which is the source of the implicit monotonic translocating assumption.

Contrary to the conventional assumption of monotonic translocation, the raw current data suggests that the motor protein drives RNA both forward and backward during sequencing. Figure 2B illustrates this with several example fragments of DRS raw current signal (21), where each fragment roughly corresponds to three neighboring 5-mers. The raw current signals, as shown in Figure 2B, strongly support this hypothesis, with several instances of measured current intensities matching both the previous and next 5-mer’s baseline. These repeated patterns, observed across multiple DRS datasets, provide empirical evidence of the jiggling translocation. Similar patterns are widely observed across the whole data. This suggests that the RNA molecule may move forward and backward while passing through the pore. This observation is also supported by previous reports (24,25), in which the helicase (the motor protein) translocates the DNA strand through the nanopore in a back-and-forth manner. Depending on ATP or ADP binding, the motor protein may translocate the DNA/RNA forward or backward by 0.5~1 nucleotides.

**Figure 2.**
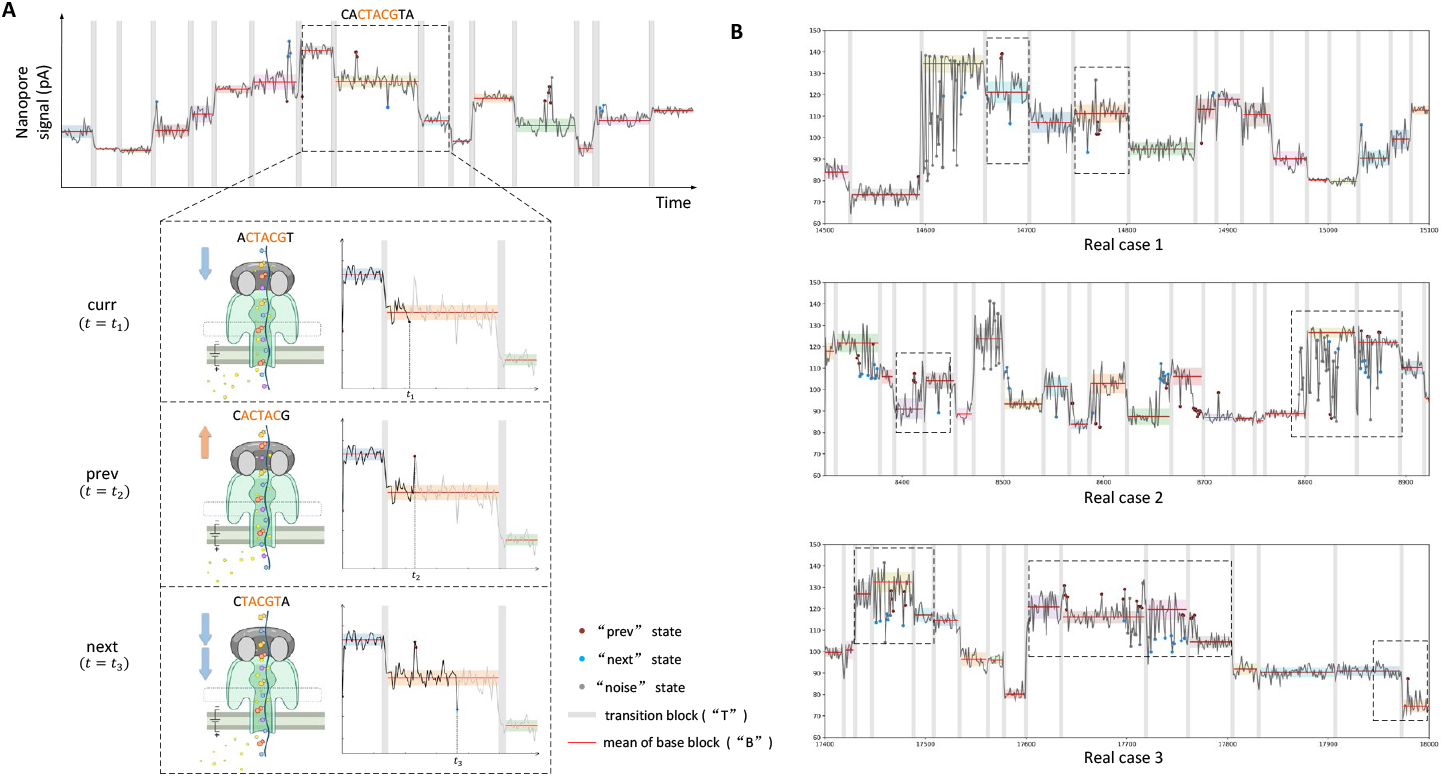
RNA translocation hypothesis. (A) Jiggling RNA translocation hypothesis. The top panel shows the raw current signal of Nanopore direct RNA sequencing, with gray areas representing SegPore-estimated transition blocks. We focus on three neighboring 5-mers, considering the central 5-mer (CTACG) as the current 5-mer. The RNA molecule may briefly move forward or backward during the translocation of the current 5-mer. If the RNA molecule is pulled backward, the previous 5-mer is placed in the pore, and the current signal (“prev” state, red dots) resembles the previous 5-mer’s baseline (mean and standard deviation highlighted by red lines and shades). If the RNA is pushed forward, the current signal (“next” state, blue dots) is similar to the next 5-mer’s baseline. (B) Example raw current signals supporting the jiggling hypothesis. The dashed rectangles highlight base blocks, with red and blue points representing measurements corresponding to the previous and next 5-mer, respectively. Red points align closely with the previous 5-mer’s baseline, and blue points match the next 5-mer’s baseline, reinforcing the hypothesis that the RNA molecule jiggles between neighboring 5-mers. The raw current signals were extracted from mESC WT samples of the training data in the m6A benchmark experiment.

Based on the reported kinetic model (24), we hypothesize that the RNA is translocated through the pore in a jiggling manner. This “jiggling” hypothesis presents a significant departure from the traditionally accepted view of unidirectional RNA translocation and aligns better with recent evidence from studies on DNA translocation (24,25). Incorporating this hypothesis into segmentation algorithms could lead to more accurate predictions of RNA modifications by accounting for the dynamic nature of translocation. On average, the motor protein sequentially translocates 5-mers on the RNA strand forward, and each 5-mer resides in the pore for a short time. During the short period of a single 5-mer, the motor protein may swiftly drive the RNA molecule forward and backward by 0.5~1 nucleotide in the translocation process of the current 5-mer (Fig. 2A), which makes the measured current intensity occasionally similar to the previous or the next 5-mer. When the motor protein does not move the RNA molecule, we hypothesize that the RNA molecule undergoes slight thermal fluctuations, causing it to oscillate slightly within the pore and produce a stable current close to its baseline. In contrast, sharp changes in current intensity between consecutive 5-mers define transition blocks, where one 5-mer is replaced by the next.

We also assume that the raw current signal of a read can be segmented into a series of alternating base and transition blocks. In the ideal case, a base block corresponds to the base state where the 5-mer resides in the pore and jiggles between neighboring 5-mers, i.e., the current 5-mer can transiently jump to the previous or the next 5-mer. A transition block corresponds to the transition state between two consecutive base states where one 5-mer translocates to the next 5-mer in the pore. The current signal should be relatively flat in the base blocks, while a sharp change is expected in the transition blocks. One challenge we encountered was the overestimation of transition blocks. This may be due to a base block actually corresponding to a sub-state of a single 5-mer, rather than each base block corresponding to a full 5-mer, leading to inflated transition counts. To address this issue, SegPore’s alignment algorithm was refined to merge multiple base blocks (which may represent sub-states of the same 5-mer) into a single 5-mer, thereby facilitating further analysis.

### SegPore Workflow

The SegPore workflow (Fig. 1) consists of five key steps: (1) Preprocess fast5 files to pair raw current signal segments with corresponding RNA sequence fragments; (2) Segment each raw current signal using a hierarchical hidden Markov model (HHMM) into base and transition blocks; (3) Align the derived base blocks with the paired RNA sequence; (4) Fit a two-component GMM to estimate the modification state at the single-molecule level or the modification rate at the site level; (5) Use the results from Step (4) to update relevant parameters. Steps (3) to (5) are iterative and continue until the estimated parameters stabilize.

The final outputs of SegPore are the events and modification state predictions. SegPore’s events are similar to the outputs of Nanopolish’s “eventalign” command, in that they pair raw current signal segments with the corresponding RNA reference 5-mers. Each 5-mer is associated with various features, such as start and end positions, mean current, and standard deviation, derived from the paired signal segment. For 5-mers that exhibit one clearly unmodified component and one clearly modified component, SegPore reports the modification rate at each site, as well as the modification state of that site on individual reads.

A key component of SegPore is the 5-mer parameter table, which specifies the mean and standard deviation for each 5-mer in both modified and unmodified states (Fig. 1A). Since the peaks (representing modified and unmodified states) are separable for only a subset of 5-mers, SegPore can provide modification parameters for these specific 5-mers. For other 5-mers, modification state predictions are unavailable.

### Segmentation benchmark

To evaluate SegPore’s performance in raw signal segmentation, we compared SegPore with Nanopolish (v0.14.0) and Tombo (v1.5.1) using three Nanopore direct RNA sequencing (DRS) datasets: two HEK293T datasets (wild type and Mettl3 knockout) (21) and the HCT116 dataset (26). Nanopolish and SegPore employed the “eventalign” method to align 5-mers on each read with their corresponding raw signals, producing the mean and standard deviation (std) of the aligned signal segments. Tombo used the “resquiggle” method to segment the raw signals, but the resulting signals are not reported on the absolute pA scale. To ensure a fair comparison with SegPore, we standardized the segments using the poly(A) tail in the same way as SegPore (See preprocessing section in Materials and Methods).

To benchmark segmentation performance, we used two key metrics (details provided in Supplementary Note 1, Section 8): (1) the log-likelihood of the segment mean, which measures how closely the segment matches ONT’s 5-mer parameter table (used as ground truth), and (2) the standard deviation (std) of the segment, where a lower std indicates reduced noise and better segmentation quality. If the raw signal segment aligns correctly with the corresponding 5-mer, its mean should closely match ONT’s reference, yielding a high log-likelihood. A lower std of the segment reflects less noise and better performance overall.

As shown in Table 1, SegPore consistently achieved the best performance averaged on all 5-mers across all datasets, with the highest log-likelihood and the lowest std values. These results suggest that SegPore provides a more accurate segmentation of the raw signal with significantly reduced noise compared to Nanopolish and Tombo. It is worth noting that the data points corresponding to the transition state between two consecutive 5-mers are not included in the calculation of the standard deviation in SegPore’s results in Table 1. However, their exclusion does not affect the overall conclusion, as there are on average only ~6 points per event in the transition state (see Supplementary Table S1 for more details).

**Table 1.** Segmentation benchmark on RNA002 data.

| Test Dataset | Avg. std ($\hat{\sigma}$) ↓ | | | Avg. log p ($\hat{L}$) ↑ | | |
| --- | --- | --- | --- | --- | --- | --- |
|  | Nanopolish | Tombo | SegPore | Nanopolish | Tombo | SegPore |
| HEK293T(WT) | 3.073 | 4.187 | **2.736** | -2.871 | -3.749 | **-2.778** |
| HEK293T(KO) | 2.948 | 4.204 | **2.670** | -2.856 | -3.749 | **-2.745** |
| HCT116 | 3.167 | 4.076 | **2.872** | -2.872 | -3.704 | **-2.746** |

To provide a more intuitive comparison, the segmentation results for two example raw signal clips are illustrated in Supplementary Fig. S1. These examples demonstrate the clearer, more precise segmentation achieved by SegPore compared to Nanopolish.

To evaluate SegPore’s performance on RNA004 data, we compared it with f5c (v1.5) (27) and Uncalled4 (28) (v4.1.0) using three public DRS datasets: the S. cerevisiae dataset (29), the curlcake IVT and m6A datasets (30). The RNA002 data provides reference current levels for 5-mers, whereas RNA004 provides reference values for 9-mers, with Uncalled 4 normalizing them to approximately zero mean and unit variance. As there are currently no established poly(A) detection methods available for RNA004, we used f5c to standardize the raw signals of each read prior to segmentation. SegPore was then applied to perform segmentation on the standardized signals. We computed the same benchmarking metrics—average log-likelihood and standard deviation—using the standardized raw signals and the segmentation results from f5c, Uncalled4, and SegPore. The 9-mer parameter table in pA scale for RNA004 data provided by f5c (see Materials and Methods) was used in the analysis. As shown in Table 2, SegPore achieved the best overall performance across all datasets, indicating its robustness and suitability for RNA004 data. Moreover, we find that the jiggling hypothesis remains valid for RNA004, as illustrated in Supplementary Fig. S4.

**Table 2.** Segmentation benchmark on RNA004 data.

| Test Dataset | Avg. std ($\hat{\sigma}$) ↓ | | | Avg. log p ($\hat{L}$) ↑ | | |
| --- | --- | --- | --- | --- | --- | --- |
|  | f5c | Uncalled4 | SegPore | f5c | Uncalled4 | SegPore |
| *S. cerevisiae* | 3.129 | 3.970 | **3.125** | -2.541 | -3.835 | **-2.489** |
| curlcake(IVT) | 3.424 | 4.729 | **3.359** | -2.716 | -2.904 | **-2.515** |
| curlcake(m6A) | 3.520 | 4.971 | **3.392** | -3.118 | -3.524 | **-2.599** |

### m6A identification at the site level

We evaluated SegPore’s performance in raw signal segmentation and m6A identification using independent public datasets as both training and test data. Since m6A modifications typically occur at DRACH motifs (where D denotes A, G, or U, and H denotes A, C, or U) (13), this study focuses on estimating m6A modifications on these motifs.

To begin, we estimated the 5-mer parameter table for m6A modifications using Nanopore direct RNA sequencing (DRS) data from three wild-type human HEK293T cell samples (21). The fast5 files from all samples were concatenated, and the full SegPore workflow was run to obtain the 5-mer parameter table. The parameter estimation process was iterated five times to ensure stabilization, refining the modification parameters for ten 5-mers where the modification state distribution significantly differs from the unmodified state.

Next, we applied SegPore’s segmentation and m6A identification to test data from wild-type mouse embryonic stem cells (mESCs) (23). Given the comparable methods and input data requirements, we benchmarked SegPore against several baseline tools, including Tombo, MINES (31), Nanom6A (32), m6Anet, Epinano (33), and CHEUI (20). By default, MINES and Nanom6A use eventalign results generated by Tombo, while m6Anet, Epinano, and CHEUI rely on eventalign results produced by Nanopolish. In Fig. 3C, “Nanopolish+m6Anet” refers to the default m6Anet pipeline, whereas “SegPore+m6Anet” denotes a configuration in which Nanopolish’s eventalign results are replaced with those from SegPore. Based on the output of SegPore eventalign, we fine-tune m6Anet using the HCT116 data at all DRACH motifs, aiming to demonstrate that the performance of SegPore benefits downstream models. Additionally, due to the differences in the availability of ground truth labels for specific k-mer motifs between human and mouse (Supplementary Fig. S2), six shared 5-mers were selected to demonstrate SegPore’s performance in modification prediction directly. By utilizing the 5-mer parameter table derived from the training data, SegPore employs a two-component GMM to calculate the modification rates at the selected m6A sites.

**Figure 3.**
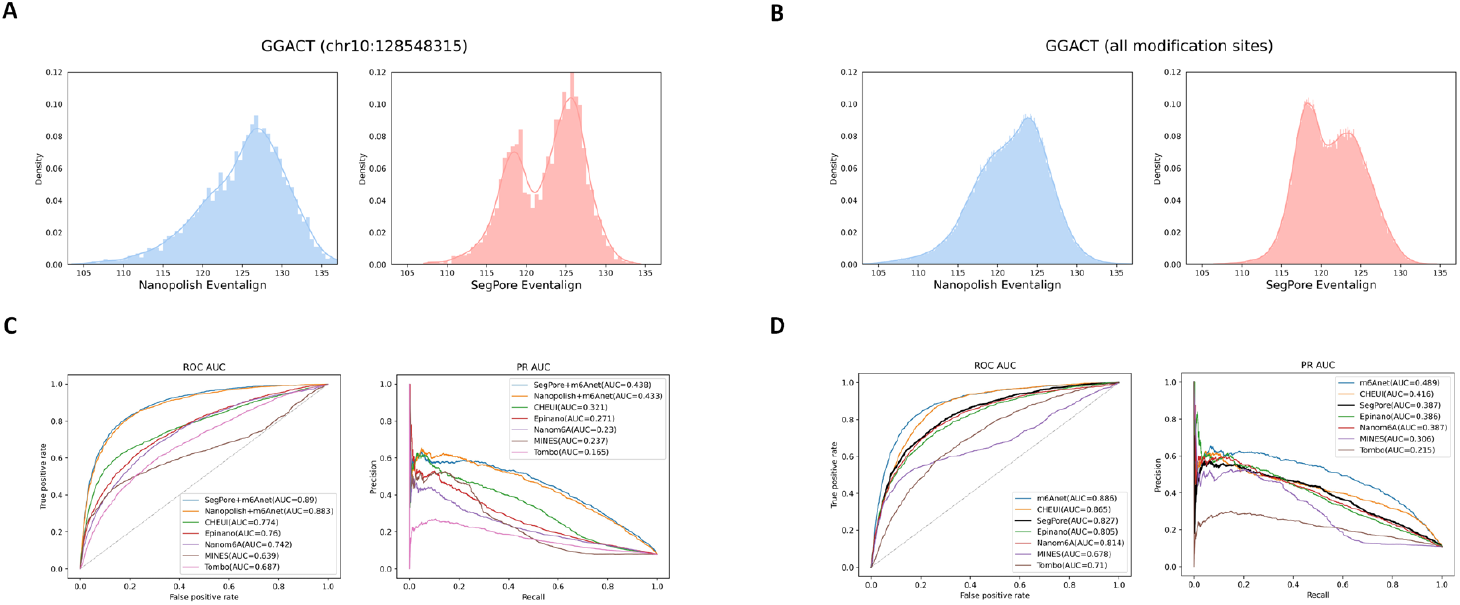
m6A identification at the site level. (A) Histogram of the estimated mean from current signals mapped to an example m6A-modified genomic location (chr10:128548315, GGACT) across all reads in the training data, comparing Nanopolish (left) and SegPore (right). The x-axis represents current in picoamperes (pA). (B) Histogram of the estimated mean from current signals mapped to the GGACT motif at all annotated m6A-modified genomic locations in the training data, again comparing Nanopolish (left) and SegPore (right). The x-axis represents current in picoamperes (pA). (C) Site-level benchmark results for m6A identification across all DRACH motifs, showing performance comparisons between SegPore+m6Anet and other methods. (D) Benchmark results for m6A identification on six selected motifs at the site level, comparing SegPore and other baseline methods.

SegPore demonstrated improved segmentation compared to Nanopolish. Figure 3A shows the eventalign results at an example genomic location with m6A modifications. SegPore’s results show a more pronounced bimodal distribution in the raw signal segment mean, indicating clearer separation of modified and unmodified signals. Furthermore, when pooling all reads mapped to m6A-modified locations at the GGACT motif, SegPore exhibited more distinct peaks (Fig. 3B), indicating reduced noise and potentially enabling more reliable modification detection.

We evaluated m6A predictions using two approaches: (1) SegPore’s segmentation results were fed into m6Anet, referred to as SegPore+m6Anet, which works for all DRACH motifs and (2) direct m6A predictions from SegPore’s Gaussian Mixture Model (GMM), which is limited to the six selected 5-mers shown in Supplementary Fig. S2C that exhibit clearly separable modified and unmodified components in the GMM (see Materials and Methods for details).

In terms of m6A identification, SegPore performed strongly on the test data. Using miCLIP2 (34) data as the ground truth, we calculated the area under the curve (AUC) for both the receiver operating characteristic (ROC) and precision-recall (PR) curves. SegPore+m6Anet achieved the best performance with an ROC AUC of 89.0% and a PR AUC of 43.8% (Fig. 3C). For six selected m6A motifs, SegPore achieved an ROC AUC of 82.7% and a PR AUC of 38.7%, earning the third best performance compared with deep leaning methods m6Anet and CHEUI (Fig. 3D). It is noteworthy that SegPore’s GMM for m6A estimation is a very simple model, utilizing far fewer parameters than DNN-based methods. Achieving the decent performance with such a simple model is a significant accomplishment. These results highlight SegPore’s robust performance in m6A identification. For practical applications, we recommend taking the intersection of m6A sites predicted by SegPore and m6Anet to obtain high-confidence modification sites, while still benefiting from the interpretability provided by SegPore’s predictions.

### m6A identification at the single molecule level

SegPore naturally identifies m6A modifications at the single-molecule level, which is crucial for understanding the heterogeneity of RNA modifications across individual transcripts. We benchmarked SegPore against CHEUI using an *in vitro* transcription (IVT) dataset containing two samples—one transcribed with m6A and the other with adenine (22). This dataset provides clear ground truth for m6A modifications at the single-molecule level, with all adenosine positions replaced by m6A in the ivt_m6A sample, and all adenosines unmodified in the ivt_normalA sample. We used 60% of the data for training and 40% for testing, with both SegPore and CHEUI estimating the m6A modification probability at each adenosine site on each read. Based on these probabilities, we calculated the ROC-AUC and PR-AUC by varying the modification probability threshold.

As shown in Figure 4A, SegPore outperformed CHEUI on this benchmark dataset, achieving better performance in both PR-AUC (94.7% vs 90.3%) and ROC-AUC (93% vs 89.9%). These results clearly demonstrate SegPore’s accuracy and robustness in detecting single-molecule m6A modifications.

**Figure 4.**
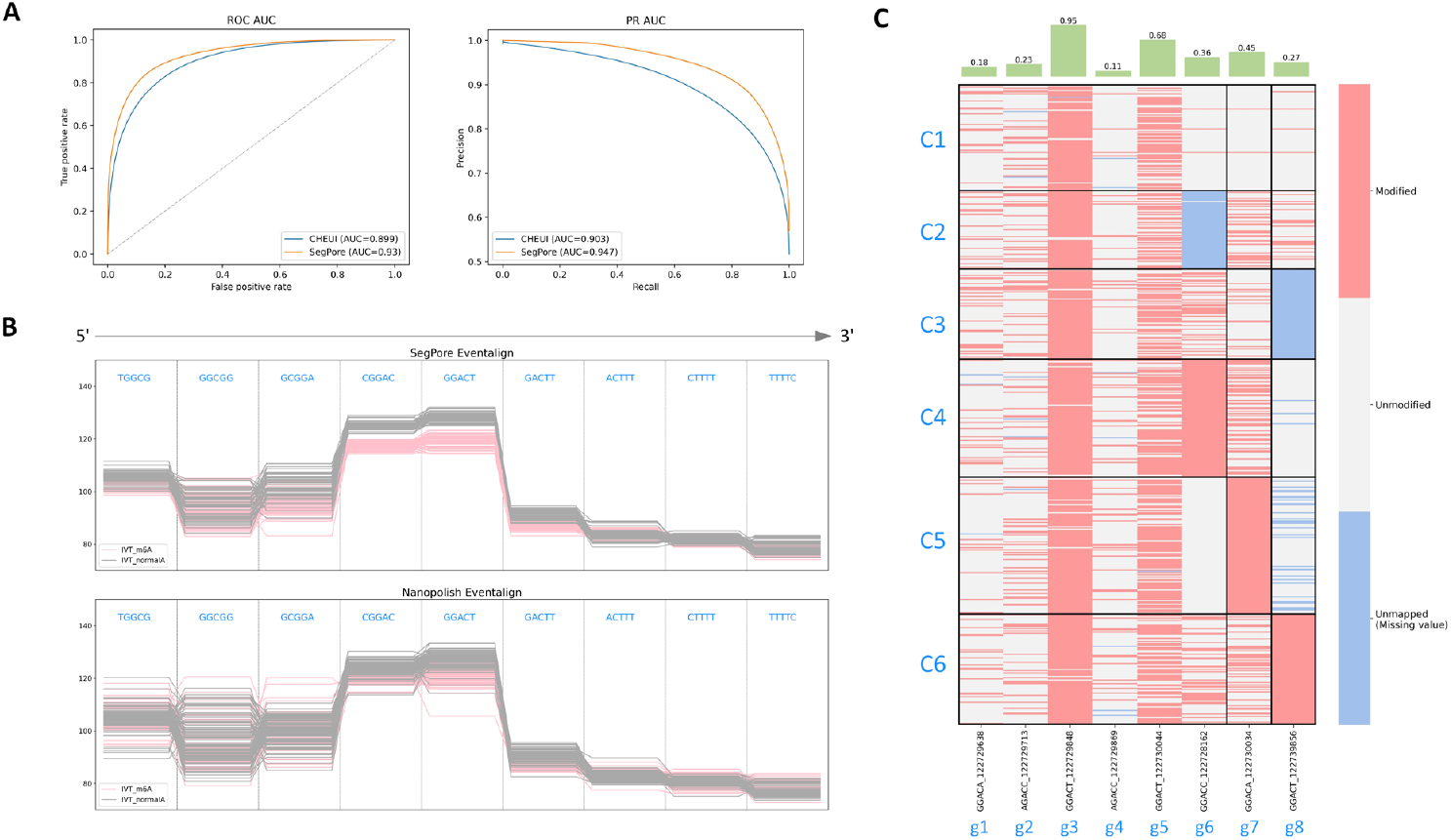
m6A identification at the single-molecule level. (A) Benchmark results for single-molecule m6A identification on IVT data. SegPore shows better performance compared to CHEUI in both PR-AUC and ROC-AUC. (B) Comparison of “eventalign” results from SegPore and Nanopolish for five consecutive k-mers. Note that DRS is sequenced from 3’ to 5’, so the k-mers enters the pore from right to left. A total of 100 reads were randomly sampled from transcript locations A1 (positions 711-719) in both the IVT_normalA and IVT_m6A samples (SRA: SRP166020). Each line represents an individual read, and the y-axis shows the raw signal intensity in picoampere (pA). Pink lines represent the IVT_m6A sample, and gray lines represent the IVT_normalA sample. The k-mers “GCGGA,” “CGGAC,” “GGACT,” “GACTT,” and “ACTTT” all contain N6-Methyladenosine (m6A) in the IVT_m6A sample. SegPore’s results show clearer separation between m6A and adenosine, especially for “CGGAC” and “GGACT,” compared to Nanopolish. (C) The upper panel shows the modification rate for selected genomic locations in the example gene ENSMUSG00000003153. The lower panel displays the modification states of all reads mapped to this gene. The black borders in the heatmap highlight the biclustering results, showing distinct modification patterns across different read clusters labeled C1 through C6.

Next, we demonstrate SegPore’s interpretability through an example comparison of raw signal clips from both ivt_m6A and ivt_normalA samples (Fig. 4B). The raw signal segments (means) are aligned to a randomly selected m6A site as well as its neighbouring sites in the reference sequence. Each line represents an individual read, with pink lines from the ivt_m6A sample and gray lines from the ivt_normalA sample. SegPore’s segmentation clearly distinguishes between m6A and adenosine at the single-molecule level, while Nanopolish’s segmentation shows less distinction.

Interestingly, we observed that the position of m6A within a 5-mer can affect the signal intensity. For instance, a clear difference between m6A and adenosine is evident when m6A occupies the fourth position in the 5-mer (e.g., “CGGAC”), but this difference is less pronounced when m6A is in the second position (e.g., “GACTT”).

To illustrate the benefits of single-molecule m6A estimation, we present an example gene (ENSMUSG00000003153) from the mESC dataset used in site-level m6A identification. Figure 4C shows a heatmap of highly modified genomic locations (modification rate > 0.1), where rows represent reads and columns represent genomic locations. Biclustering reveals six clusters of reads and three clusters of genomic locations.

The heatmap in Figure 4C suggests heterogeneity in m6A modification patterns across different reads of the same gene. Biclustering reveals that modifications at g6 are specific to cluster C4, g7 to cluster C5, and g8 to cluster C6, while the first five genomic locations (g1 to g5) show similar modification patterns across all reads. Additionally, high modification rates are observed at the 3rd and 5th positions across the majority of reads. These results suggest that m6A modification patterns can vary significantly even within a single gene. This observation highlights the complexity of RNA modification regulation and underscores the importance of single-molecule resolution for understanding RNA function at a finer scale.

## DISCUSSION

One of the main computational challenges in direct RNA sequencing (DRS) lies in accurately segmenting the raw current signal. We developed a segmentation algorithm that models the jiggling property in the physical process of DRS, resulting in cleaner current signals for m6A identification at both the site and single-molecule levels. Our results demonstrate that SegPore’s segmentation enables clear differentiation between m6A-modified and unmodified adenosines. We believe that the de-noised current signals will be beneficial for other downstream tasks, such as the estimation of m5C, pseudouridine, and other RNA modifications. Nevertheless, several open questions remain for future research. In SegPore, we assume a drastic change between two consecutive 5-mers, which may hold for 5-mers with large difference in their current baselines but may not hold for those with small difference. As with other RNA modification estimation methods, SegPore can be affected by misalignment errors, particularly when the baseline signals of adjacent k-mers are similar. These cases may lead to spurious bimodal signal distributions and require careful interpretation. Another key question concerns the physical interpretation of the derived base blocks. Ideally, one base block would correspond to a single 5-mer, but in practice, multiple base blocks often align with one 5-mer. We hypothesize that the HHMM may segment a 5-mer’s current signal into multiple base blocks, where the 5-mer oscillates between different sub-states, each characterized by distinct baselines.

Currently, SegPore models only the modification state of the central nucleotide within the 5-mer. However, modifications at other positions may also affect the signal, as shown in Figure 4B. Therefore, introducing multiple states for a 5-mer to account for modifications at multiple positions within the same 5-mer could help to improve the performance of the model. This approach, however, would significantly increase model complexity—introducing two states per location results in 2^5^ = 32 modification states per 5-mer, a challenge for simple GMMs. It remains to be investigated how to model multiple modification states properly to improve the performance in RNA modification estimation. While the current two-state assumption simplifies the model, it has proven effective, as demonstrated by improved performance on m6A benchmarks at both site and single-molecule levels.

A key advantage of SegPore is its interpretability, which sets it apart from DNN-based methods. SegPore provides a clearer understanding of RNA modification predictions. A limitation of DNN-based approaches is that they often struggle to differentiate between m6A and adenosine signals, as their intensity levels are quite similar (Fig. 4B, Supplementary Fig. S3). This poses a challenge when applying DNN-based methods to new datasets without short read sequencing-based ground truth. In such cases, it is difficult for researchers to confidently determine whether a predicted m6A modification is genuine. SegPore encodes current intensity levels for different 5-mers in a parameter table, where unmodified and modified 5-mers are modeled using two Gaussian distributions. One can generally observe a clear difference in the intensity levels between 5-mers with an m6A and those with adenosine, which makes it easier for a researcher to interpret whether a predicted m6A site is genuine (see Supplementary Fig. S5). A challenge is the relatively small number of 5-mers that show significant changes in their modification states. To improve accuracy, larger training datasets and expanding the scope to 7-mers or 9-mers could help capture more context, potentially revealing more significant baseline changes.

The limited improvement of SegPore combined with m6Anet over Nanopolish+m6Anet in bulk *in vivo* analysis (Fig. 3) may be explained by several factors: potential alignment inaccuracies due to pseudogenes or transcript isoforms, the complexity of *in vivo* datasets containing additional RNA modifications (e.g., m5C, m7G) affecting signal baselines, and the fact that m6Anet is specifically trained on events produced by Nanopolish rather than SegPore. Additionally, the lack of a modification-free control (*in vitro* transcribed) sample in the

benchmark dataset makes it difficult to establish true baselines for each k-mer. Despite these limitations, SegPore demonstrates clear improvement in single-molecule m6A identification in IVT data (Fig. 4), suggesting it is particularly well suited for i*n vitro* transcription data analysis.

Although SegPore provides clear interpretability, there is potential to explore DNN-based models that can directly leverage SegPore’s segmentation results. Currently, m6Anet computes features (e.g., mean and standard deviation) from raw signal segments, which are then fed into a neural network for m6A prediction. However, a more direct approach—where raw signal segments are used as input to a DNN—could allow for the extraction of more intricate features that may exist within the signal but are currently underexplored. These high-order features might capture subtle aspects of the raw signal, leading to improved m6A estimation. Since SegPore currently models only the mean and standard deviation, further work involving advanced DNNs could extend beyond this, uncovering finer patterns in the signal that traditional statistical models might miss.

Computation speed is also a concern when processing fast5 files. We addressed this by implementing a GPU-accelerated inference algorithm in SegPore, resulting in a 10-to 20-fold speedup compared to the CPU-based version. We believe that the GPU-implementation will unlock the full potential of SegPore for a wider range of downstream tasks and larger datasets. SegPore’s running times on datasets of varying sizes, using a single NVIDIA DGX-1 V100 GPU and one CPU core, are provided in Supplementary Fig. S6.

In summary, we developed a novel software SegPore that considered the conformation changes of motor protein to segment raw current signal of Nanopore direct RNA sequencing. SegPore effectively masks out noise in the raw signal, leading to improved m6A identification at both site and single-molecule levels.

## DATA AVAILABILITY

The data utilized in this study are obtained from publicly available repositories. Details regarding the accession number and data processing can be found in Methods. The resulting data and the source code are hosted on GitHub (https://github.com/guangzhaocs/SegPore).

## AUTHOR CONTRIBUTIONS

Guangzhao Cheng: Formal analysis, Methodology, Validation, Writing—original draft & editing. Aki Vehtari: Writing—review. Lu Cheng: Conceptualization, Formal analysis, Methodology, Writing—original draft & review.

## ACKNOWLEDGEMENTS

We would like to thank Prof. Zhijie Tan from Wuhan University for a useful discussion about the molecule dynamics of Nanopore sequencing, Dr. Dan Zhang from Sichuan University for helpful tutorials about Nanopore analysis workflows. We also thank Prof. Luo Guanzheng for sharing the m6A benchmark baseline results. G.C. and L.C. acknowledge the computational resources provided by the Aalto Science-IT project.

