## Supplementary Figures, Tables, and Notes for "Raw signal segmentation for estimating RNA modification from Nanopore direct RNA sequencing data"

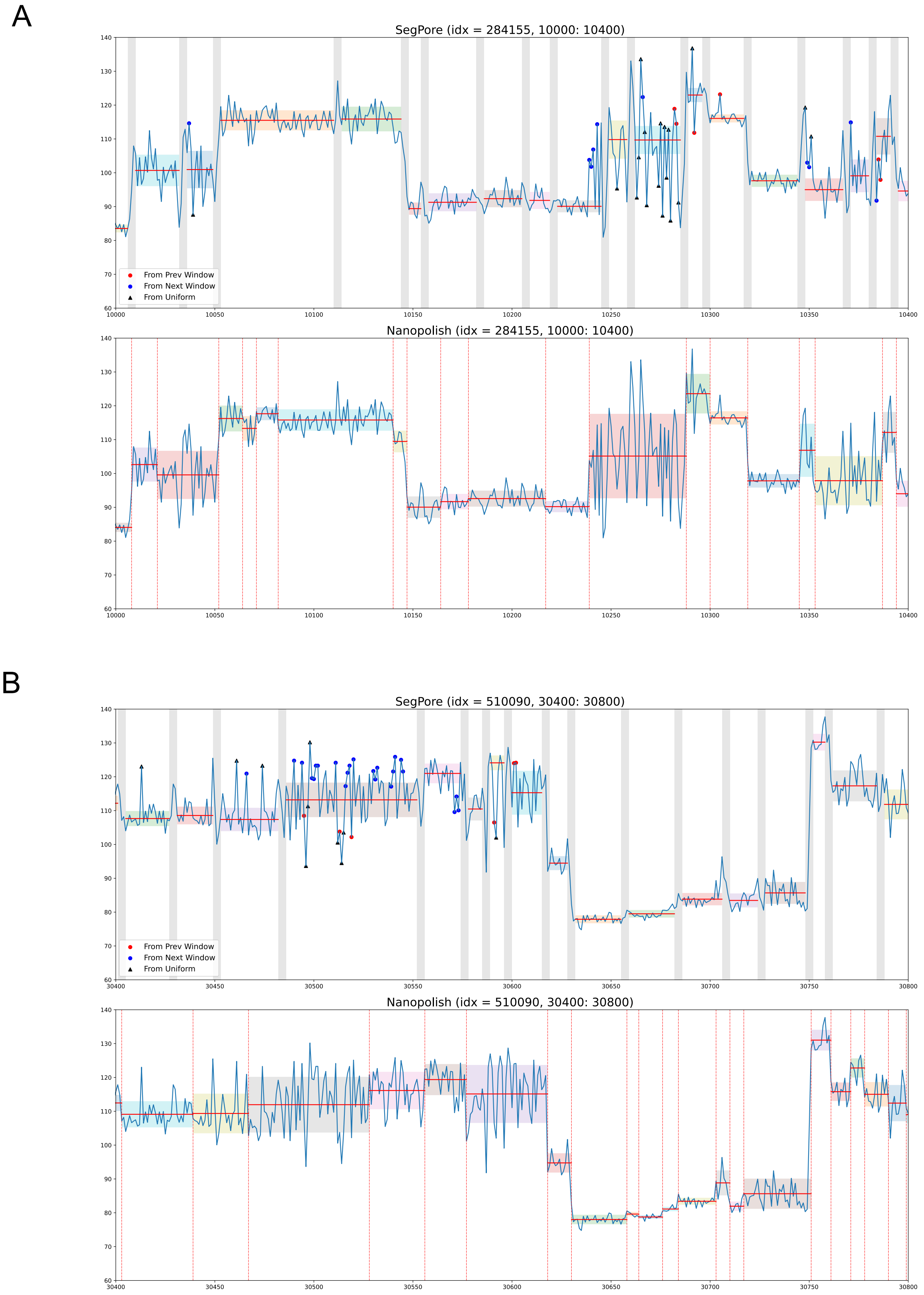

**Supplementary Figure S1.** Comparison of raw signal segmentation results between SegPore and Nanopolish on RNA002 data. The same raw signal clip is used for both

SegPore (top panel) and Nanopolish (bottom panel). Two example raw signal clips (*A*, *B*) are taken from the mES_WT sample (SRA: SRP166020) in the m6A identification section. The x-axis represents time, and the y-axis represents the raw current signal intensity (pA). The vertical bars or lines indicate the borders between segments (transition blocks). The horizontal shaded areas represent the estimated standard deviation of each base block, and the red horizontal line indicates the mean of each base block. There are four states in the base blocks: “prev” (red dot), “next” (blue dot), “curr,” and “noise” (black triangle).

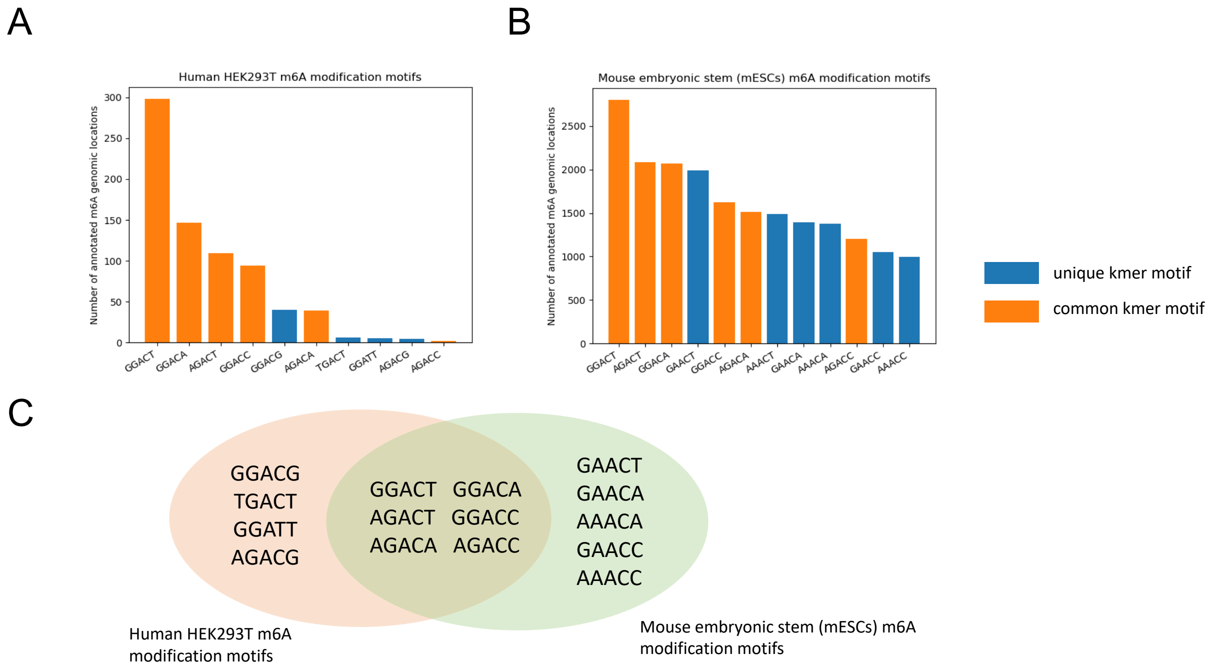

**Supplementary Figure S2.** m6A k-mer statistics differences between the human HEK293T (training data) and the mouse mES_WT (test data). (*A*) Number of modification sites for each selected 5-mer (in total 10 motif 5-mers) based on the ground truth of the human data (ENA: PRJEB40872). (*B*) Number of modification sites for all 5-mers (in total 11 motif 5-mers) based on the ground truth of the mouse data (SRA: SRP166020). The ground truth is obtained by taking the union of MeRIP-seq, miCLIP, and miCLIP2. (*C*) Venn diagram of the two 5-mer motif sets (training data and test data).

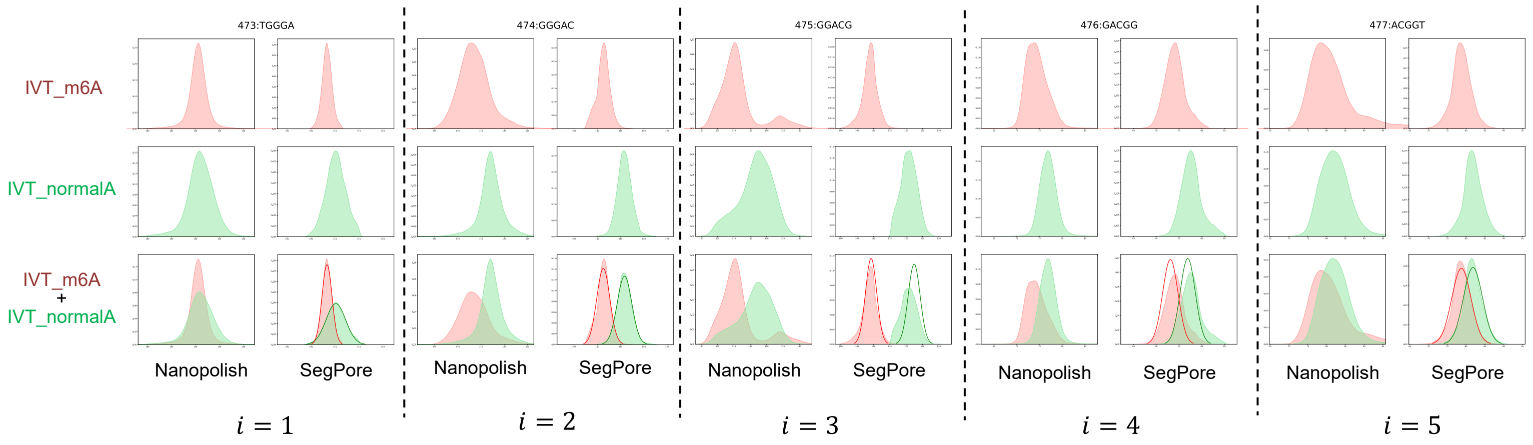

**Supplementary Figure S3**. Comparison of eventalign of SegPore and Nanopolish on consecutive 5-mers. The figures show the distribution of estimated segment means for each k-mer in transcript locations A2 (position 473-477) in IVT_m6A and IVT_normalA samples (SRA: SRP166020). All reads aligned to these 5-mers are used to generate the marginal distribution. The third row is the overlay of the distributions of IVT_m6A and IVT_normalA samples. m6A are located at different positions of the given 5-mer. It can be seen that the peaks are relatively well separated for the third and fourth 5-mers in SegPore's result, but not so clear in Nanopolish's result.

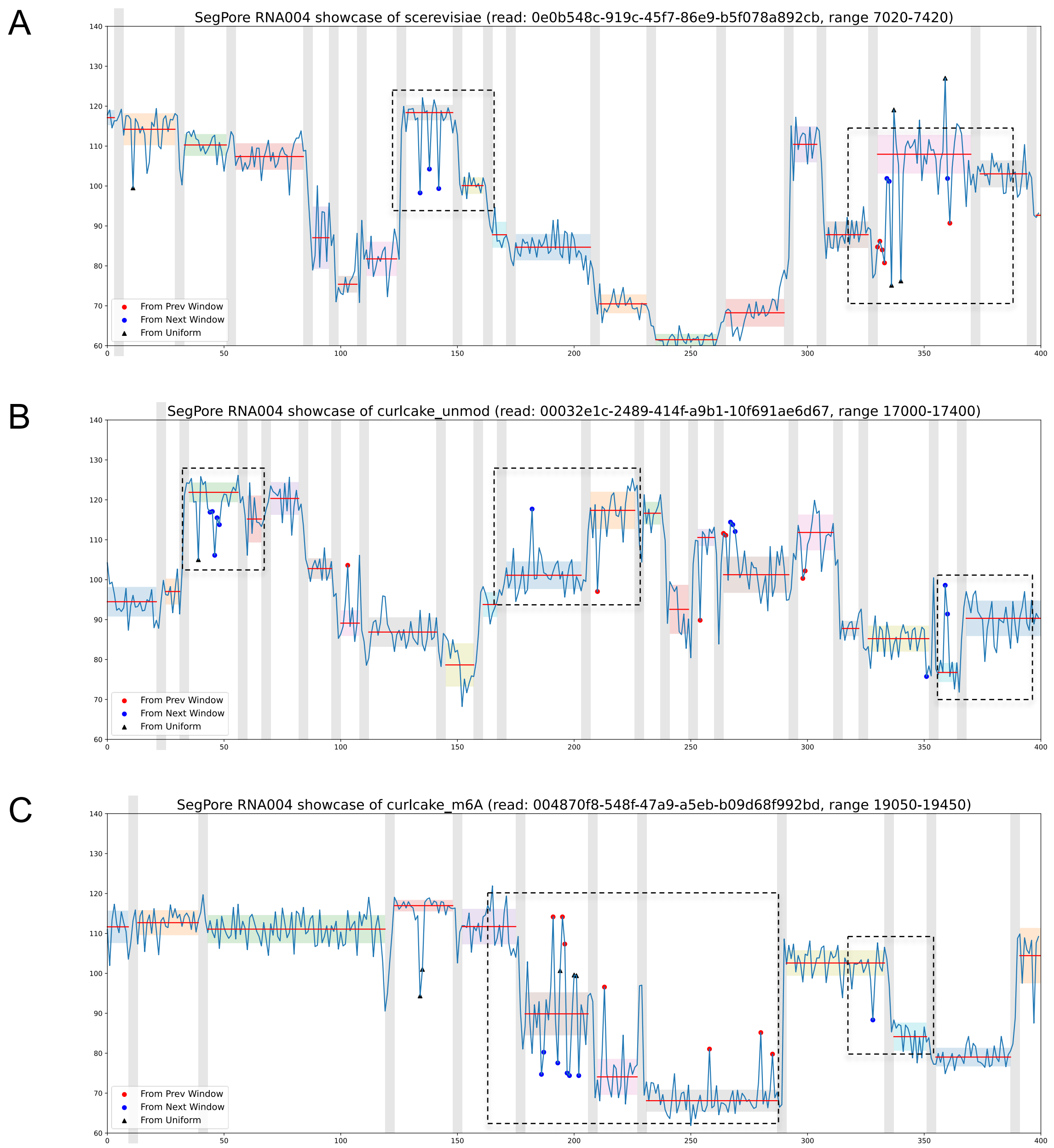

**Supplementary Figure S4.** RNA translocation hypothesis illustrated on RNA004 data. (*A*) Example from the S. cerevisiae sample (SRA: SRP527927). (*B*) Example from the curlcake_unmod sample (PRJEB82528). (*C*) Example from the curlcake_m6A sample (PRJEB82528). In all panels, the x-axis shows time, and the y-axis shows raw current signal intensity (pA, transformed using f5c scale parameters). Vertical lines mark the boundaries between segments (transition blocks). Horizontal shaded areas indicate the estimated standard deviation for each base block, while the red horizontal line shows the mean signal level for that block. Each base block is annotated with four possible states: “prev” (red dot), “next” (blue dot), “curr,” and “noise” (black triangle).

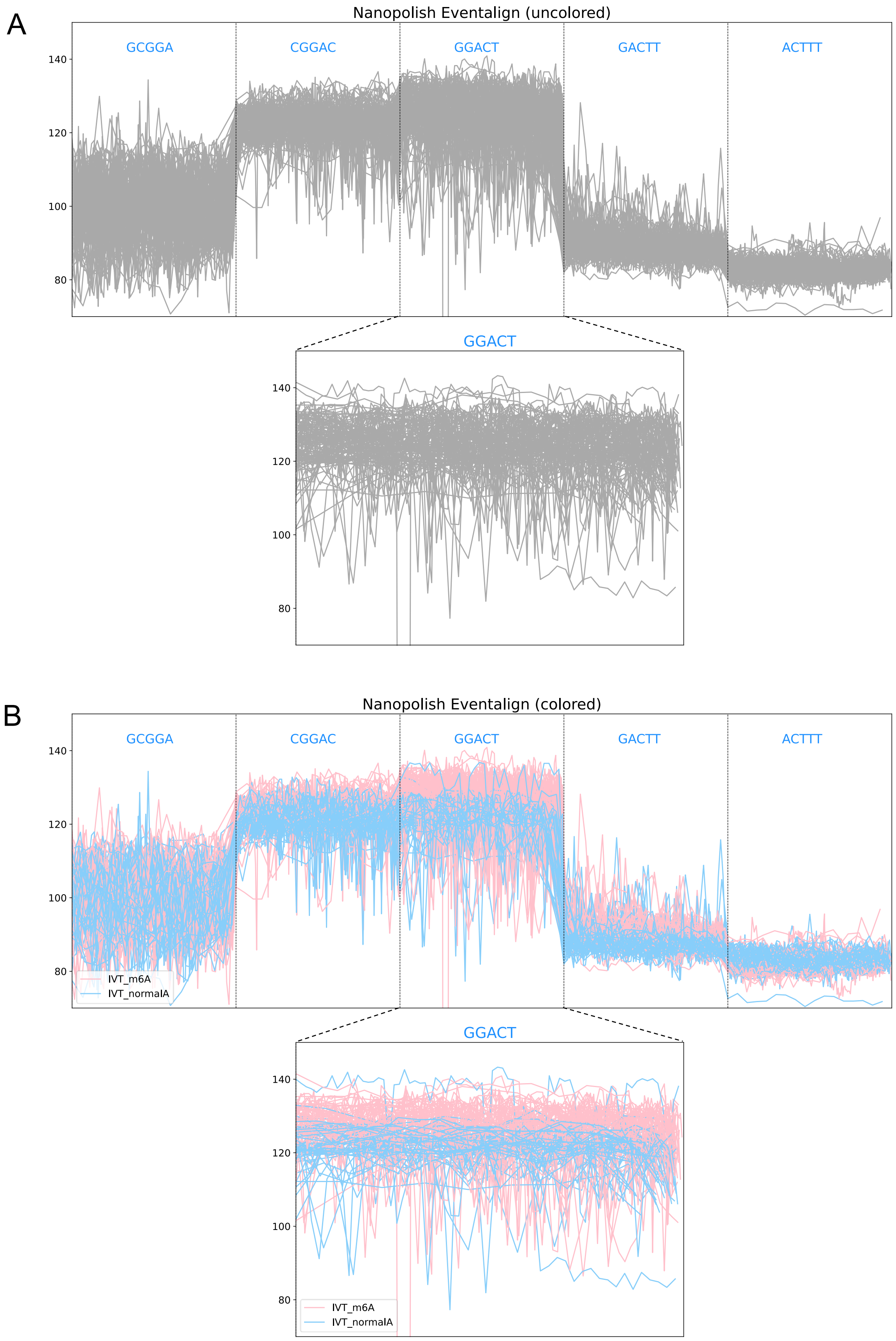
A
Nanopolish Eventalign (uncolored)
GCGGA
CGGAC
GGACT
GACTT
ACTTT
140
120
100
80
GGACT
140
120
100
80
B
Nanopolish Eventalign (colored)
GCGGA
CGGAC
GGACT
GACTT
ACTTT
140
120
100
80
IVT_m6A
IVT_normalA
GGACT
140
120
100
80
IVT_m6A
IVT_normalA

**Supplementary Figure S5.** Illustration of Nanopolish raw signal segmentation with eventalign results. This figure corresponds to Figure 4B (Nanopolish Eventalign) in the manuscript. (*A*) Visualization without color differentiation: IVT_m6A and IVT_normalA samples are shown in the same color. (*B*) Visualization with color differentiation: IVT_m6A and IVT_normalA samples are shown in distinct colors for clarity. In both panels, the top section displays five consecutive k-mers, while the bottom section provides a zoomed-in view of the central GGACT k-mer. Without color contrast, baseline differences in the GGACT signals between m6A and normalA samples are not easily detectable, as the baselines appear closely overlapping. Furthermore, identifying other subtle signal features that differentiate the two GGACT states—features leveraged by deep neural networks—remains challenging for field researchers.

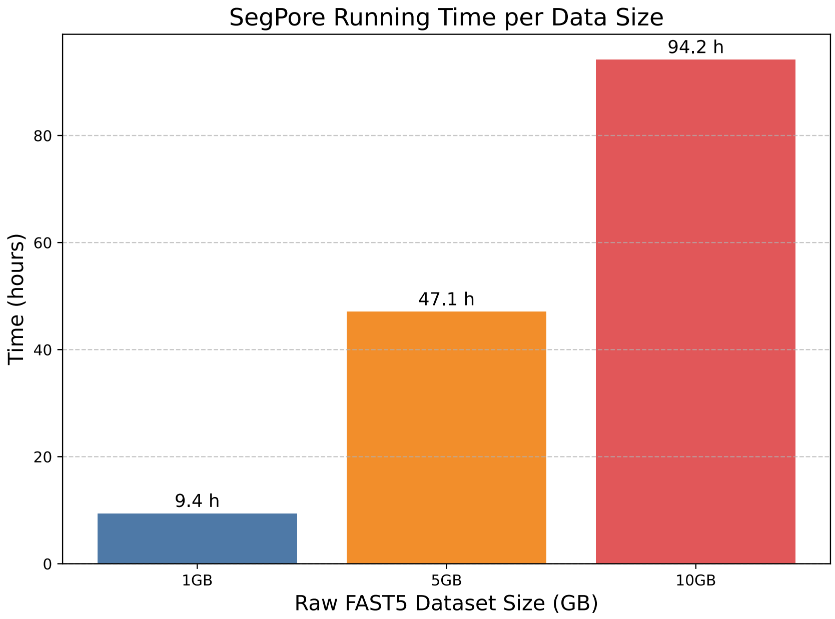

**Supplementary Figure S6.** Running time of SegPore on datasets of varying sizes. The figure shows the total duration required for the segmentation task using a single NVIDIA DGX-1 V100 GPU and one CPU core with 32 GB system memory allocated.

**Supplementary Table S1.** Segmentation benchmark on the RNA002 dataset excluding boundary regions. This table corresponds to Table 1 in the main text. For Nanopolish and Tombo, the first and last three points of each event were removed prior to recalculating the metrics.

| Test Dataset | Avg. std ($\hat{\sigma}$) ↓ | | | Avg. log p ($\hat{L}$) ↑ | | |
|---|---|---|---|---|---|---|
| | Nanopolish | Tombo | SegPore | Nanopolish | Tombo | SegPore |
| HEK293T(WT) | 2.862 | 4.090 | **2.736** | -2.838 | -3.772 | **-2.778** |
| HEK293T(KO) | 2.856 | 4.120 | **2.670** | -2.878 | -3.729 | **-2.745** |
| HCT116 | 3.058 | 4.053 | **2.872** | -2.869 | -3.708 | **-2.746** |

### SegPore workflow

(Supplementary Note 1)

Guangzhao Cheng, Aki Vehtari, Lu Cheng

October, 2025

#### Contents

**1 Notation** **2**

**2 Introduction** **2**

**3 Basecalling, mapping and preprocessing** **4**

**4 Hierarchical hidden Markov model** **4**

**7 Estimation of $k$-mer parameters table** **12**

**8 Evaluation of segmentation and event alignment** **13**

### 1 Notation

| | |
|---|---|
| $\lvert\cdot\rvert$ | Take the length of the inside vector. |
| $\boldsymbol{Y}$ | All reads in the input multi-fast5 or pod5 file. $\boldsymbol{Y} = (\boldsymbol{y_1}, \boldsymbol{y_2}, \cdots, \boldsymbol{y_i}, \cdots)$. $\boldsymbol{y_i}$ are the raw current measurements of $i$th read. Assuming there are $N$ reads, i.e. $\lvert\boldsymbol{Y}\rvert = N$. |
| $\boldsymbol{y_i}$ | The current signal of $i$th read. $\boldsymbol{y_i} = (y_{i1}, y_{i2}, \cdots, y_{ij}, \cdots, y_{il_i})$, where $l_i$ is the length of $\boldsymbol{y_i}$, i.e. $\lvert\boldsymbol{y_i}\rvert = l_i$ . |
| $\boldsymbol{y_i}^{(k)}$ | The $k$th mapped current signal fragment of $i$th read $\boldsymbol{y_i}$, given by eventalign using Nanopolish. |
| $\boldsymbol{R}$ | Reference sequences for each read. $\boldsymbol{R} = (\boldsymbol{r_1}, \boldsymbol{r_2}, \cdots, \boldsymbol{r_i}, \cdots)$. |
| $\boldsymbol{r_i}$ | The reference sequence corresponding to $i$th read $\boldsymbol{y_i}$, given by basecalling and minimap2. $\boldsymbol{r_i} = (r_{i1}, r_{i2}, \cdots, r_{ij}, \cdots, r_{im_i})$, where $r_{ij} \in \{A, C, G, T\}$ and $m_i$ is the length of $\boldsymbol{r_i}$, i.e. $\lvert\boldsymbol{r_i}\rvert = m_i$. |
| $\boldsymbol{r_i}^{(k)}$ | The $k$th reference sequence fragment of $\boldsymbol{r_i}$ corresponding to $\boldsymbol{y_i}^{(k)}$. Note that $\lvert\boldsymbol{y_i}^{(k)}\rvert$ is larger than $\lvert\boldsymbol{r_i}^{(k)}\rvert$ in general. |
| $\boldsymbol{s_i}$ | The $k$-mer list corresponds to $\boldsymbol{r_i}$. We have $\boldsymbol{s_i} = (s_{i1}, s_{i2}, \cdots, s_{ij}, \cdots, s_{im_i})$, where $s_{ij} = r_{i,j-2}r_{i,j-1}r_{ij}r_{i,j+1}r_{i,j+2}$ and $s_{ij} \in \{AAAAA, AAAAC, AAAAG, AAAAT, \cdots, TTTTT\}$. Note that $\lvert\boldsymbol{s_i}\rvert = \lvert\boldsymbol{r_i}\rvert = m_i$. |
| $\boldsymbol{s_i}^{(k)}$ | The $k$-mer list fragment corresponds to the $k$th reference sequence fragment $\boldsymbol{r_i}^{(k)}$. $\lvert\boldsymbol{s_i}^{(k)}\rvert = \lvert\boldsymbol{r_i}^{(k)}\rvert$. |
| $\boldsymbol{Z}$ | The estimated states of all reads given by the Hierarchical Hidden Markov Model (HHMM). $\boldsymbol{Z} = (\boldsymbol{z_1}, \boldsymbol{z_2}, \cdots, \boldsymbol{z_i}, \cdots)$. |
| $\boldsymbol{z_i}$ | The hidden states for each current measure of $i$th read. $\boldsymbol{z_i} = (z_{i1}, z_{i2}, \cdots, z_{ij}, \cdots, z_{il_i})$, where $z_{ij} \in \{0, 1, 2, 3, 4\}$ corresponds to $y_{ij}$ and $\lvert\boldsymbol{y_i}\rvert = \lvert\boldsymbol{z_i}\rvert = l_i$, where $z_{ij} = 0$ means the transition state and $z_{ij} = 1, 2, 3, 4$ means base states. |
| $\boldsymbol{z_i}^{(k)}$ | The $k$th fragment in $\boldsymbol{z_i}$, which corresponds to $\boldsymbol{y_i}^{(k)}$. Note that $\lvert\boldsymbol{z_i}^{(k)}\rvert = \lvert\boldsymbol{y_i}^{(k)}\rvert$. |
| $\boldsymbol{\mu_i}$ | The mean values of base segments given by HHMM, which are estimated from $\boldsymbol{y_i}$ and $\boldsymbol{z_i}$. $\boldsymbol{\mu_i} = (\mu_{i1}, \mu_{i2}, \cdots, \mu_{ij}, \cdots, \mu_{in_i})$, where $j = 1 \cdots n_i$ indexes the base segments of $i$th read. In general, we have $l_i > n_i > m_i$. |
| $\boldsymbol{\mu_i}^{(k)}$ | The $k$th fragment of $\boldsymbol{\mu_i}$, which corresponds to $\boldsymbol{y_i}^{(k)}$ and $\boldsymbol{z_i}^{(k)}$. $\boldsymbol{\mu_i}^{(k)} = (\mu_{i1}^{(k)}, \mu_{i2}^{(k)}, \cdots, \mu_{ij}^{(k)}, \cdots)$. In general we have $\lvert\boldsymbol{y_i}^{(k)}\rvert > \lvert\boldsymbol{\mu_i}^{(k)}\rvert > \lvert\boldsymbol{s_i}^{(k)}\rvert$. |
| $\boldsymbol{u_i}^{(k)}$ | The modification state of each $k$-mer in $\boldsymbol{s_i}^{(k)}$. $\boldsymbol{u_i}^{(k)} = (u_{i1}^{(k)}, u_{i2}^{(k)}, \cdots, u_{ij}^{(k)}, \cdots)$, where $u_{ij}^{(k)} \in \{un, mod\}$ indicates if $s_{ij}^{(k)}$ is in unmodified or modified state. $\lvert\boldsymbol{u_i}^{(k)}\rvert = \lvert\boldsymbol{s_i}^{(k)}\rvert$. |
| $\boldsymbol{T}^{ref}$ | The reference $k$-mer (5-mer) parameter table provided by Oxford Nanopore Technology (ONT). $\boldsymbol{T}^{ref} = \{\ (\mu_{s,u}, \sigma_{s,u}),\ \forall s \in \{AAAAA, AAAAC, \cdots, TTTTT\}, u = \text{“}un\text{”}\ \}$. |
| $\boldsymbol{T}^{kmer}$ | The parameter table for all 5-mers. $\boldsymbol{T}^{kmer} = \{\ (\mu_{s,u}, \sigma_{s,u}),\ \forall s \in \{AAAAA, AAAAC, \cdots, TTTTT\}, \forall u \in \{\text{“}un\text{”}, \text{“}mod\text{”}\}\ \}$. |

### 2 Introduction

Given the parameter table $\boldsymbol{T}^{kmer}$ for all $k$-mers, the overall workflow of SegPore consists of the following major steps: (1) Base-calling and mapping, (2) preprocessing of nanopore raw signal, (3) segmenting nanopore raw signal using hierarchical hidden Markov model (HHMM), and (4) aligning raw signal segments with corresponding $k$-mer list. The input of Segpore is the raw current signal of Nanopore direct RNA sequencing. The output of Segpore workflow is the pairing of raw signal segments with corresponding $k$-mers with inferred state (unmodified state or modified state). Unless otherwise noted, the following analysis focuses on RNA002 data, using 5-mers rather than $k$-mers.

As illustrated in Fig. 1, we denote the input FAST5/POD5 file as a set of raw current signal reads $\boldsymbol{Y} = (\boldsymbol{y_1}, \boldsymbol{y_2}, \cdots, \boldsymbol{y_i}, \cdots)$. We use $\lvert\boldsymbol{Y}\rvert$ to denote the number of reads and $\lvert\boldsymbol{y_i}\rvert$ to denote the number of measurements / data points of $\boldsymbol{y_i}$. We then perform basecalling using Guppy and map basecalled sequence to the reference sequences (Sec. 3), yielding a pairing between the $i$th read $\boldsymbol{y_i}$ and its corresponding reference sequence $\boldsymbol{r_i}$. We assume there are $k$ matched fragments $[(\boldsymbol{y_i}^{(1)}, \boldsymbol{r_i}^{(1)}), (\boldsymbol{y_i}^{(2)}, \boldsymbol{r_i}^{(2)}), \cdots, (\boldsymbol{y_i}^{(k)}, \boldsymbol{r_i}^{(k)})]$. For the $j$th fragment $\boldsymbol{y_i}^{(j)}$, we use HHMM (Sec. 4) to estimate the hidden states for each measurement $\boldsymbol{z_i}^{(j)} = (z_{i1}^{(j)}, z_{i2}^{(j)}, \cdots)$, where $z_{im}^{(j)} \in \{0, 1, 2, 3, 4\}$ and $\lvert\boldsymbol{y_i}^{(j)}\rvert = \lvert\boldsymbol{z_i}^{(j)}\rvert$. Based on $\boldsymbol{y_i}^{(j)}$ and $\boldsymbol{z_i}^{(j)}$, we can convert it into a list of low variation segments, with their means denoted by $\boldsymbol{\mu_i}^{(j)} = (\mu_{i1}^{(j)}, \mu_{i2}^{(j)}, \cdots)$. As a result, the number of measurements in $\boldsymbol{y_i}^{(j)}$ is

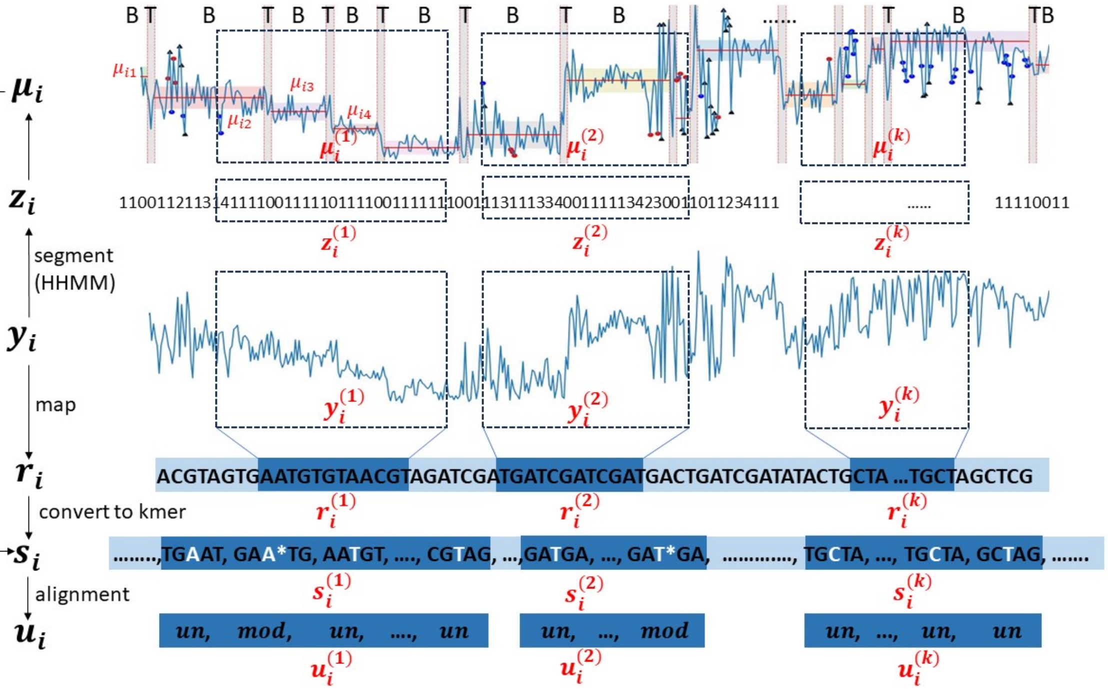

Figure 1: SegPore workflow with a fixed 5-mer parameter table $\boldsymbol{T}^{kmer}$. The input is the raw current signal $\boldsymbol{y_i}$ and reference sequence $\boldsymbol{r_i}$ for $i$th read. The output is the paired current signal segments $\boldsymbol{\mu_i}$, 5-mer list $\boldsymbol{s_i}$ and modification states $\boldsymbol{u_i}$.

| modle_kmer | un_mu | un_sigma | mod_mu | mod_sigma |
|---|---|---|---|---|
| AAAAA | 108.9 | 2.68 | - | - |
| AAAAC | 107.75 | 2.68 | - | - |
| AAAAG | 101.72 | 2.68 | - | - |
| AAAAT | 112.77 | 2.68 | - | - |
| AAACA | 99.38 | 2.41 | 95.28 | 2.44 |
| AAACC | 100 | 1.32 | 96.82 | 2.42 |
| AAACG | 101.01 | 3.45 | - | - |
| AAACT | 106.91 | 2.18 | 101.98 | 2.29 |
| AAAGA | 110.54 | 4.06 | - | - |
| AAAGC | 107.69 | 4.06 | - | - |
| AAAGG | 108.29 | 4.06 | - | - |
| AAAGT | 108.73 | 4.06 | - | - |
| AAATA | 114.11 | 3.11 | - | - |
| ...... | ...... | ...... | ...... | ...... |
| TTTTT | 80.78 | 1.97 | - | - |

Figure 2: Example of $\boldsymbol{T}^{kmer}$ (1024 × 4). “-” represents “ NA”.

larger than the number of segments in $\boldsymbol{\mu}_{\boldsymbol{i}}^{(j)}$, i.e. $|\boldsymbol{y}_{\boldsymbol{i}}^{(j)}| = |\boldsymbol{z}_{\boldsymbol{i}}^{(j)}| > |\boldsymbol{\mu}_{\boldsymbol{i}}^{(j)}|$. After that we perform full or partial alignment between the raw signal segments $\boldsymbol{\mu}_{\boldsymbol{i}}^{(j)}$ and reference sequence $\boldsymbol{r}_{\boldsymbol{i}}^{(j)}$, which is converted into a list of 5-mers $\boldsymbol{s}_{\boldsymbol{i}}^{(j)} = (s_{i1}^{(j)}, s_{i2}^{(j)}, \cdots)$ and $s_{im}^{(j)} =$ "$r_{i,m-2}^{(j)} r_{i,m-1}^{(j)} r_{i,m}^{(j)} r_{i,m+1}^{(j)} r_{i,m+2}^{(j)}$". Based on $\boldsymbol{\mu}_{\boldsymbol{i}}^{(j)}$ and $\boldsymbol{s}_{\boldsymbol{i}}^{(j)}$, we perform full or partial alignment (Sec. 5) to match each 5-mer with one or multiple current signal segments, as well as estimating its modification state $\boldsymbol{u}_{\boldsymbol{i}}^{(j)} = (u_{i1}^{(j)}, u_{i2}^{(j)}, \cdots)$, where $u_{im}^{(j)} \in \{$"*un*", "*mod*"$\}$.

However, $\boldsymbol{T}^{kmer}$ is not fully known beforehand (Fig. 2), especially the parameters for the modification states of 5-mers. We have to estimate the parameters from the training data through an iterative process (Sec. 7). After obtaining the estimated $\boldsymbol{T}^{kmer}$, we derive the modification states from new test data using SegPore workflow with fixed $\boldsymbol{T}^{kmer}$.

### 3 Basecalling, mapping and preprocessing

This section describes the details about the basecalling and mapping process , as well as raw data preprocessing for standardizing the raw signal of all reads.

First, we obtain the matching between raw current signals and reference sequences. The raw multi-fast5 reads (input file) were basecalled using Guppy (v6.0.1)[1], which results in a FASTQ file. Then we align the yielded FASTQ file to the reference sequences (we use cDNA reference) using minimap2 (v2.24-r1122)[2], which provides a bam file that matches the basecalled reads with the reference sequence. After that we feed the raw fast5 file, basecalled FASTQ file, bam file and reference sequence to Nanopolish (v0.13.2)[3] to get (1) matching fragments between the raw current signal $\boldsymbol{y}_{\boldsymbol{i}}^{(k)}$ and the reference sequence $\boldsymbol{r}_{\boldsymbol{i}}^{(k)}$ (as well as $\boldsymbol{s}_{\boldsymbol{i}}^{(k)}$) and (2) the current signal fragment for poly(A) part of the mRNA molecule.

Then, we standardize the raw current signal based on the poly(A) tail. Due to technical reasons, there exist variations in the current signal of the same sequence fragment across different reads. It is necessary to perform a standardization. Since all reads have a poly(A) tail in Nanopore direct RNA sequencing, we could standardize the reads such that the mean and std of poly(A) part is the same for all reads. The standardization is given by

$$standardized\_signal = \frac{raw\_signal - \mu_{polyA}}{\sigma_{polyA}} * \sigma_{standPolyA} + \mu_{standPolyA} \quad (1)$$

where $\mu_{standPolyA} = 108.9$ and $\sigma_{standPolyA} = 1.67$ (RNA002). Note that $\boldsymbol{y}_{\boldsymbol{i}}$ refers to the standardized signal throughout this document.

### 4 Hierarchical hidden Markov model

This section describes the hierarchical hidden Markov model (HHMM) to segment the raw current signal of a read. The input of HHMM is $\boldsymbol{y}_{\boldsymbol{i}}$ and the output is $\boldsymbol{z}_{\boldsymbol{i}}$. For notation simplicity, we use $\boldsymbol{y}$ and $\boldsymbol{z}$ to represent $\boldsymbol{y}_{\boldsymbol{i}}$ and $\boldsymbol{z}_{\boldsymbol{i}}$ in this section, respectively.

#### 4.1 Model assumptions

Here we have several assumptions regarding the raw signal. We first assume there is a sharp change of the baseline levels of the current signal between two consecutive 5-mers. This means there is a large change of current signal within a short period of time. If we fit a line to the corresponding signals, we expect to see a large slope. The second assumption is that the 5-mer resides in the Nanopore oscillates forward and backward during a 5-mer event, while staying in the pore most of the time. Based on these two assumptions, we can divide the raw current signal into two states (hidden states of the outer HMM): the base states for 5-mer event and the transition state for the transition between two consecutive 5-mer events. We assume that the current signal always starts with a base state, then

[1] https://community.nanoporetech.com/docs/prepare/library_prep_protocols/Guppy-protocol/v/gpb_2003_v1_revax_14dec2018/guppy-software-overview

[2] https://github.com/lh3/minimap2

[3] https://nanopolish.readthedocs.io/en/latest/index.html

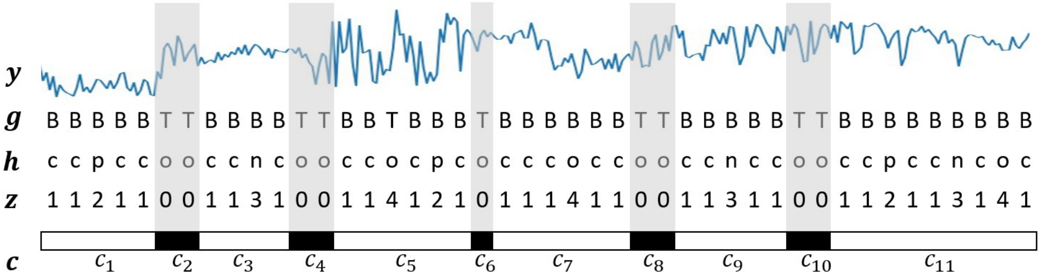

Figure 3: Example of HHMM where $g_i \in \{\text{“}B\text{”}, \text{“}T\text{”}\}$, $h_i \in \{\text{“}curr\text{”}, \text{“}prev\text{”}, \text{“}next\text{”}, \text{“}noise\text{”}\}$ and $c_i \in \{\text{“}B\text{”}, \text{“}T\text{”}\}$. In this figure, “c”, “p”, “n” and “o” represent “curr”, “prev”, “next” and “noise” respectively in $\boldsymbol{h}$. $|\boldsymbol{y}| = |\boldsymbol{g}| = |\boldsymbol{h}|$ and $|\boldsymbol{c}| = 11$.

switches to a transition state, after that switches back to a base state or transition state alternatively, in the end the signal should end in a base state. For the base state, we use an inner HMM to model the oscillations, which has four hidden states:“curr”, “prev”, “next” and “noise”. For the transition state, we use a linear regression model to model the abrupt change of the current signal.

#### 4.2 Outer HMM

We define the following notations for the HHMM. As illustrated in Fig. 3, we assume there are $N$ time points of the current signal, i.e. $\boldsymbol{y} = (y_1, y_2, \cdots, y_i, \cdots, y_N)$. We use $\boldsymbol{g} = (g_1, g_2, \cdots, g_i, \cdots, g_N), g_i \in \{\text{“}B\text{”}, \text{“}T\text{”}\}$ to indicate the hidden states of the outer HMM, where $B$ and $T$ stand for “base” and “transition”, respectively. Similarly, we use $\boldsymbol{h} = (h_1, h_2, \cdots, h_i, \cdots, h_N), h_i \in \{\text{“}curr\text{”}, \text{“}prev\text{”}, \text{“}next\text{”}, \text{“}noise\text{”}\}$, respectively. We then integrate $\boldsymbol{g}$ and $\boldsymbol{h}$ into a single vector $\boldsymbol{z} = (z_1, z_2, \cdots, z_i, \cdots, z_N)$, where $z_i = \tilde{g}_i \cdot \tilde{h}_i$. Here $\tilde{g}_i$ (“B”→1, “T”→0) and $\tilde{h}_i$ (“curr”→1, “prev”→2, “next”→3, “noise”→4) are the numerical mappings of $g_i$ and $h_i$. Based on $\boldsymbol{g}$, we can derive that $\boldsymbol{y}$ is divided into $2K+1$ blocks, which is defined by $\boldsymbol{c} = (c_1, c_2, \cdots, c_i, \cdots, c_{2K+1}), c_i \in \{\text{“}B\text{”}, \text{“}T\text{”}\}$. Note that $c_i = \text{“}B\text{”}\ \forall i = 1, 3, \cdots, 2K+1$ and $c_i = \text{“}T\text{”}\ \forall i = 2, 4, \cdots, 2K$. As a result, we could also represent $\boldsymbol{y} = (\boldsymbol{y}^{(1)}, \boldsymbol{y}^{(2)}, \cdots, \boldsymbol{y}^{(i)}, \cdots, \boldsymbol{y}^{(2K+1)})$, where $\boldsymbol{y}^{(i)}$ represent the $i$th block. The base blocks are given by $(\boldsymbol{y}^{(b_1)}, \boldsymbol{y}^{(b_2)}, \cdots, \boldsymbol{y}^{(b_i)}, \cdots, \boldsymbol{y}^{(b_{K+1})})$, where $b_i = 2i-1$ and $i \in \{1, 2, \cdots, K+1\}$. The transition blocks are given by $(\boldsymbol{y}^{(t_1)}, \boldsymbol{y}^{(t_2)}, \cdots, \boldsymbol{y}^{(t_i)}, \cdots, \boldsymbol{y}^{(t_K)})$, where $t_i = 2i$ and $i \in \{1, 2, \cdots, K\}$. Similarly, we have $\boldsymbol{g} = (\boldsymbol{g}^{(1)}, \boldsymbol{g}^{(2)}, \cdots, \boldsymbol{g}^{(i)}, \cdots, \boldsymbol{g}^{(2K+1)})$ and $\boldsymbol{h} = (\boldsymbol{h}^{(1)}, \boldsymbol{h}^{(2)}, \cdots, \boldsymbol{h}^{(i)}, \cdots, \boldsymbol{h}^{(2K+1)})$.

We use $T^{outer} = \{T^{outer}_{BB}, T^{outer}_{BT}, T^{outer}_{TB}, T^{outer}_{TT}\}$ to denote the transition probabilities between the base state and the transition state in the outer HMM. We use $T^{inner} = \{T^{inner}_{h_i h_j}\ \forall h_i, h_j \in \{\text{“}curr\text{”}, \text{“}prev\text{”}, \text{“}next\text{”}, \text{“}noise\text{”}\}\}$ to denote the transition probabilities between four hidden states of the inner HMM. For base blocks, we have inner HMM parameters $\boldsymbol{\phi} = (\phi^{(b_1)}, \phi^{(b_2)}, \cdots, \phi^{(b_{K+1})})$ and $\boldsymbol{\sigma} = (\sigma^{(b_1)}, \sigma^{(b_2)}, \cdots, \sigma^{(b_{K+1})})$. For transition blocks, we have linear coefficients $\boldsymbol{\beta}_0 = (\beta_0^{(t_1)}, \beta_0^{(t_2)}, \cdots, \beta_0^{(t_K)})$, $\boldsymbol{\beta}_1 = (\beta_1^{(t_1)}, \beta_1^{(t_2)}, \cdots, \beta_1^{(t_K)})$ and noise std $\boldsymbol{\sigma}_{\boldsymbol{\epsilon}} = (\sigma_\epsilon^{(t_1)}, \sigma_\epsilon^{(t_2)}, \cdots, \sigma_\epsilon^{(t_K)})$.

The joint likelihood of the HHMM is given by

$$\begin{aligned} p(\boldsymbol{y}, \boldsymbol{g}) &= p(\boldsymbol{y}|\boldsymbol{g}) p(\boldsymbol{g}) \\ &= p(\boldsymbol{y}|\boldsymbol{g}) \pi^{outer}_{g_1} \prod_{i=2}^{N} T^{outer}_{g_{i-1} g_i} \\ &= \prod_{i=1}^{N} p(y_i|g_i) \pi^{outer}_{g_1} \prod_{i=2}^{N} T^{outer}_{g_{i-1} g_i} \end{aligned} \tag{2}$$

, where $\pi^{outer}_{g_1} = 1$ is the initial probability of hidden state of the $y_1$ (always “B” state). However, it is not possible to directly derive $p(y_i, h_i|g_i)$, since $y_i$ are not independent given $g_i$. Here we use different models for each block $c_k$ to calculate the joint probability $p(\boldsymbol{y}^{(k)}|\boldsymbol{g}^{(k)})$ to approximate $\prod_{i=1}^{N} p(y_i|g_i)$. Therefore, we have

| State | Event | Emission distribution | Paramter Estimation |
|---|---|---|---|
| "curr" | current 5-mer resides in pore | $N\left(\phi^{(b_i)}, (\sigma^{(b_i)})^2\right)$ | Yes |
| "prev" | previous 5-mer resides in pore | $N\left(\phi^{(b_{i-1})}, (\sigma^{(b_{i-1})})^2\right)$ | Yes |
| "next" | next 5-mer resides in pore | $N\left(\phi^{(b_{i+1})}, (\sigma^{(b_{i+1})})^2\right)$ | Yes |
| "noise" | unknown noise | $Unif(lb, ub)$ | No |

Table 1: Hidden states of inner HMM for the $b_i$th block.

$$
\begin{aligned}
\prod_{i=1}^{N} p(y_i|g_i) &= \prod_{j=0}^{2K+1} p\Big(\boldsymbol{y}^{(j)} \mid c_j\Big) \\
&= \prod_{j=0}^{K} p\Big(\boldsymbol{y}^{(2j+1)} \mid c_{2j+1} = \text{“}B\text{”}\Big) \prod_{j=1}^{K} p\Big(\boldsymbol{y}^{(2j)} \mid c_{2j} = \text{“}T\text{”}\Big)
\end{aligned} \tag{3}
$$

The first part is the marginal likelihood for base blocks, while the second part is the marginal likelihood for transition blocks. The specific likelihood formulas are given in the following sections.

For the outer HMM, the transition probabilities $T^{outer}$ are pre-specified and the emission probabilities are obtained from the inner HMM (Sec. 4.3) and linear regression model (Sec. 4.4). The initial probability for the first hidden state $\pi^{outer}_{g_1}$ is set to 1, since the first state should always be "B".

#### 4.3 Inner HMM for base state

This section provides the derivations for calculating the likelihood of the $i$th base block $b_i$, where $i \in \{1, 2, \cdots, K+1\}$. We assume the data $\boldsymbol{y}^{(b_i)} = (y_1^{(b_i)}, y_2^{(b_i)}, \cdots, y_{N_{b_i}}^{(b_i)})$ is generated by an HMM with four states: "curr", "prev", "next", and "noise". The descriptions of the states are given by Table 1, where parameters for the noise state are given and parameters for other states need to be estimated from the data.

The likelihood for $k$th block is given by

$$
\begin{aligned}
& p\Big(\boldsymbol{y}^{(b_i)} \mid c_{b_i} = \text{“}B\text{”}\Big) = \sum_{\boldsymbol{h}^{(b_i)}} p\Big(\boldsymbol{y}^{(b_i)}, \boldsymbol{h}^{(b_i)} \mid c_{b_i} = \text{“}B\text{”}\Big) \\
&= \sum_{\boldsymbol{h}^{(b_i)}} p\Big(\boldsymbol{y}^{(b_i)} \mid \boldsymbol{\phi}, \boldsymbol{\sigma}, \boldsymbol{h}^{(b_i)}\ , c_{b_i} = \text{“}B\text{”}\Big) p\Big(\boldsymbol{h}^{(b_i)} \mid c_{b_i} = \text{“}B\text{”}\Big) \\
&= \sum_{\boldsymbol{h}^{(b_i)}} \left[ \prod_{j=1}^{N_{b_i}} p(y_j^{(b_i)} | h_j^{(b_i)}, \boldsymbol{\phi}, \boldsymbol{\sigma}) \pi^{inner}_{h_1^{(b_i)}} \prod_{j=2}^{N_{b_i}} T^{inner}_{h_{j-1}^{(b_i)} h_j^{(b_i)}} \right] \\
&= \sum_{\boldsymbol{h}^{(b_i)}} \Bigg[ \prod_{j=1}^{N_{b_i}} \bigg\{ N\Big(y_j^{(b_i)} | \phi^{(b_i)}, (\sigma^{(b_i)})^2\Big)^{I(h_j^{(b_i)} = \text{“}curr\text{”})} N\Big(y_j^{(b_i)} | \phi^{(b_{i-1})}, (\sigma^{(b_{i-1})})^2\Big)^{I(h_j^{(b_i)} = \text{“}prev\text{”})} \\
&\quad N\Big(y_j^{(b_i)} | \phi^{(b_{i+1})}, (\sigma^{(b_{i+1})})^2\Big)^{I(h_j^{(b_i)} = \text{“}next\text{”})} Unif\Big(y_j^{(b_i)} | lb, ub\Big)^{I(h_j^{(b_i)} = \text{“}noise\text{”})} \bigg\} \pi^{inner}_{h_1^{(b_i)}} \prod_{j=2}^{N_{b_i}} T^{inner}_{h_{j-1}^{(b_i)} h_j^{(b_i)}} \Bigg],
\end{aligned} \tag{4}
$$

where $\pi^{inner}_{h_1^{(b_i)}} = 0.25$ is the probability of the first hidden state and $I(\cdot)$ is the indicator function.

For the inner HMM, the transition probabilities $T^{inner}$ are pre-specified and the initial probability for the first hidden state $\pi^{inner}_{h_1^{(b_i)}} = 0.25$ is set to 0.25. Other parameters (see Table 1) need to be estimated.

By comparing the emission probabilities of the outer HMM (Eq. 3) and the marginal likelihood of the inner HMM (Eq. 4), we have

$$p\Big(\boldsymbol{y}^{(b_i)} \mid c_{b_i} = \text{“}B\text{”}\Big) = \prod_{j=1}^{|\boldsymbol{y}^{(b_i)}|} p(y_j^{(b_i)}|g_j^{(b_i)}) \tag{5}$$

To approximate $p(y_j^{(b_i)}|g_j^{(b_i)})$, we assume all values of $p(y_j^{(b_i)}|g_j^{(b_i)})$ are the same for any given $j$, we have

$$p\Big(\boldsymbol{y}^{(b_i)} \mid c_{b_i} = \text{“}B\text{”}\Big) = \prod_{j=1}^{|\boldsymbol{y}^{(b_i)}|} p(y_j^{(b_i)}|g_j^{(b_i)}) = [\hat{p}(y_j^{(b_i)}|g_j^{(b_i)})]^{|\boldsymbol{y}^{(b_i)}|} \tag{6}$$

The approximated emission probability of each data point in $i$th base block is given by

$$\hat{p}(y_j^{(b_i)}|g_j^{(b_i)}) = \sqrt[|\boldsymbol{y}^{(b_i)}|]{\hat{p}\Big(\boldsymbol{y}^{(b_i)} \mid c_{b_i} = \text{“}B\text{”}\Big)}, \tag{7}$$

where $\hat{p}\Big(\boldsymbol{y}^{(b_i)} \mid c_{b_i} = \text{“}B\text{”}\Big)$ can be obtained by our inference algorithm.

Note that the hidden states of the inner HMM $\boldsymbol{h}$ can be provided as a by-product of our inference algorithm. We will use it to derive the final composite hidden state $\boldsymbol{z}$ of the HHMM (Sec. 4.5), which is utilized by downstream alignment tasks (Sec. 5.3).

#### 4.4 Linear model for transition state

This section provides the derivations for calculating the likelihood of the $i$th transition block $t_i$, where $i \in \{1, 2, \cdots, K\}$. We denote the input data by $\boldsymbol{y}^{(t_i)} = (y_1^{(t_i)}, y_2^{(t_i)}, \cdots, y_{N_{t_i}}^{(t_i)})$ and its corresponding index by $\boldsymbol{x}^{(t_i)} = (1, 2, \cdots, N_{t_i})$. We assume a linear model for data points in the transition state, given by

$$\boldsymbol{y}^{(t_i)} = \beta_1^{(t_i)}\boldsymbol{x}^{(t_i)} + \beta_0^{(t_i)} + \epsilon, \tag{8}$$

where $\beta_1^{(t_i)}$ is the slope, $\beta_0^{(t_i)}$ is the intercept and $\epsilon \sim N(0, (\sigma_\epsilon^{(t_i)})^2)$ is the Gaussian noise.

The likelihood of the linear model is given by

$$\begin{aligned} p\Big(\boldsymbol{y}^{(t_i)} \mid c_{t_i} = \text{“}T\text{”}\Big) &= p\Big(\boldsymbol{y}^{(t_i)} \mid \boldsymbol{x}^{(t_i)}, \beta_0^{(t_i)}, \beta_1^{(t_i)}, \sigma_\epsilon^{(t_i)}\Big) \\ &= \prod_{j=1}^{N_{t_i}} p\Big(y_j^{(t_i)}|x_j^{(t_i)}, \beta_0^{(t_i)}, \beta_1^{(t_i)}, \sigma_\epsilon^{(t_i)}\Big) \\ &= \prod_{j=1}^{N_{t_i}} N\Big(y_j^{(t_i)} - \beta_1^{(t_i)}x_j^{(t_i)} - \beta_0^{(t_i)} \mid 0, (\sigma_\epsilon^{(t_i)})^2\Big) \end{aligned} \tag{9}$$

Similar to the inner HMM, the approximated emission probability for each data point in $i$th transition block is given by

$$\hat{p}(y_j^{(t_i)}|g_j^{(t_i)}) = \sqrt[|\boldsymbol{y}^{(t_i)}|]{\hat{p}\Big(\boldsymbol{y}^{(t_i)} \mid c_{t_i} = \text{“}T\text{”}\Big)}, \tag{10}$$

where $\hat{p}\Big(\boldsymbol{y}^{(b_i)} \mid c_{b_i} = \text{“}T\text{”}\Big)$ can be obtained by our inference algorithm.

#### 4.5 Parameter Inference

Due to the complexity of GPU-accelerate inference algorithm, the details are provided in **Supplementary Note 2**.

The inference algorithm will infer the following parameters: (1) the hidden states $\boldsymbol{g}$ of the outer HMM; (2) the hidden states of the inner HMM $\boldsymbol{h}$; (3) relevant parameters for emissions (Table 1) of each current signal fragment of the $2k+1$th block (base block) $\boldsymbol{y}^{(2k+1)}$, where $k \in (0, 1, \cdots, K)$ and $c_{2k+1} = \text{“}B\text{”}$; and (4) the linear coefficients and noise std of the $2k$th block (transition block) $\boldsymbol{y}^{(2k)}$, where $k \in (1, 2, \cdots, K)$ and $c_{2k} = \text{“}T\text{”}$.

We then integrate $\boldsymbol{g}$ and $\boldsymbol{h}$ into a single vector $\boldsymbol{z} = (z_1, z_2, \cdots, z_i, \cdots, z_N)$, where $z_i = \tilde{g}_i \cdot \tilde{h}_i$ and $\tilde{g}_i, \tilde{h}_i$ are the numerical mappings of $g_i, h_i$. $\boldsymbol{z}$ is the final output of HHMM.

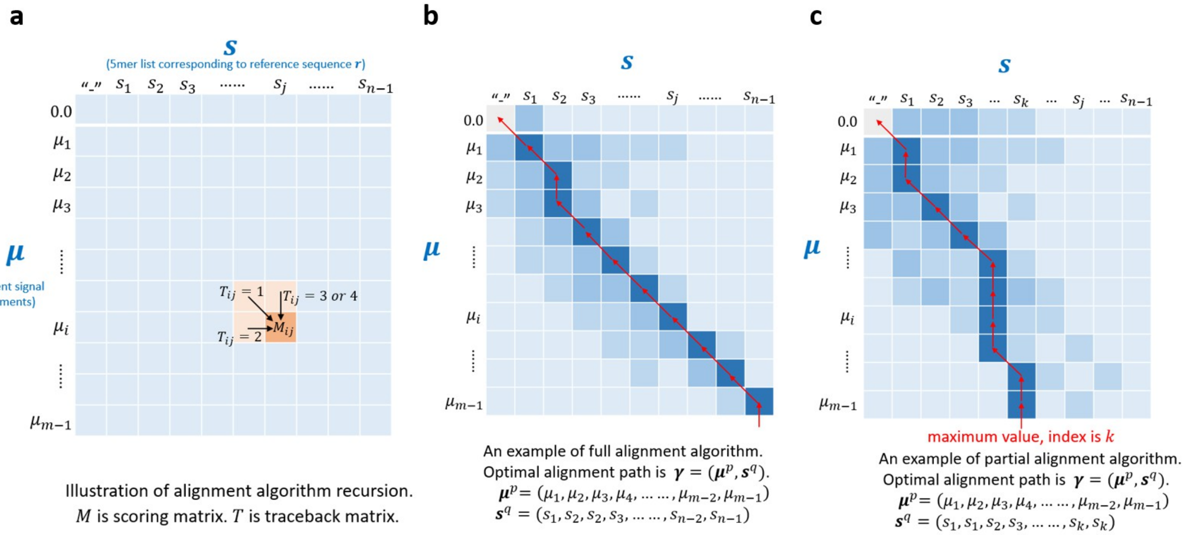

Figure 4: Alignment of signal segments with reference sequence.

### 5 Alignment of signal segments with reference sequence

This section describes the alignment algorithms for aligning an input fragment of signal segments $\boldsymbol{\mu}_{\boldsymbol{i}}^{(k)}$ and the corresponding 5-mer list $\boldsymbol{s}_{\boldsymbol{i}}^{(k)}$. After post-processing of the alignment path, we could get the modification states $\boldsymbol{u}_{\boldsymbol{i}}^{(k)}$. For notation simplicity, we drop the indeices and denote $\boldsymbol{\mu}_{\boldsymbol{i}}^{(k)}$, $\boldsymbol{s}_{\boldsymbol{i}}^{(k)}$ and $\boldsymbol{u}_{\boldsymbol{i}}^{(k)}$ by $\boldsymbol{\mu}$, $\boldsymbol{s}$ and $\boldsymbol{u}$, respectively. In general, one 5-mer $s_j$ is aligned with $k$ signal segments $(\mu_i, \mu_{i+1}, \cdots, \mu_{i+k-1})$, where $k \geq 1$.

Here we discuss different scenarios of aligning $s_j$ with $\mu_i$. The first scenario (scenario 1) is that $\mu_i$ is generated by unmodified 5-mer $s_j$, for which we could compute the likelihood using the Normal distribution defined by $(\mu_{s_j}^{un}, \sigma_{s_j}^{un}) = T_{s_j,un}^{ref}$. The second senario (scenario 2) is the $\mu_i$ is generated by modified 5-mer $s_j$, i.e. $N(\mu_{s_j}^{mod}, \sigma_{s_j}^{mod})$, where $(\mu_{s_j}^{mod}, \sigma_{s_j}^{mod}) = T_{s_j,mod}^{ref}$. The third scenario (scenario 3) is that $\mu_i$ is generated by an unknown nucleotide insertion event, whose probability is given by a pre-defined uniform distribution $p_{unif}$. The fourth scenario (scenario 4) is that the 5-mer $s_j$ represents a deletion event on the reference sequence, whose probability is given by the same uniform distribution $p_{unif}$. We define the scoring function as

$$f(s_j, \mu_i) = \begin{cases} max\Big\{log\Big(N\big(\mu_i \mid \mu_{s_j}^{un}, (\sigma_{s_j}^{un})^2\big)\Big),\ log\Big(N\big(\mu_i \mid \mu_{s_j}^{mod}, (\sigma_{s_j}^{mod})^2\big)\Big)\Big\} & (scenario\,1, 2) \\ log\big(p_{unif}\big) \quad \#s_j = \text{“}-\text{”}\ or\ \mu_i = 0.0 & (scenario\,3, 4) \end{cases} \tag{11}$$

We can modify the classical global alignment algorithm to align $\boldsymbol{\mu}$ with $\boldsymbol{s}$. The log probability of aligning $s_j$ with $\mu_i$ could be used as the matching score (Eq. 11) in the standard global alignment algorithm. Since one 5-mer $s_j$ could be aligned with multiple signal segments $(\mu_i, \mu_{i+1}, \cdots, \mu_{i+k-1})$, we need to modify the recursion such that $\mu_{i+1}$ could still be aligned with $s_j$ even if $\mu_i$ is already aligned with $s_j$. Therefore, we have the following recursions in our alignment algorithm (Fig. 4a). Here we denote the scoring matrix by $M$ $(m \times n)$ and the traceback matrix by $T$, where $|\boldsymbol{\mu}| = m - 1$ and $|\boldsymbol{s}| = n - 1$.

$$M(i, j) = max \begin{cases} M(i-1, j-1) + f(\mu_i, s_j) & \#scenario\,1, 2 \quad T(i, j) = 1 \\ M(i, j-1) + f(0.0, s_j) & \#scenario\,3 \quad T(i, j) = 2 \\ M(i-1, j) + f(\mu_i, \text{“}-\text{”}) & \#scenario\,4 \quad T(i, j) = 3 \\ M(i-1, j) + f(\mu_i, s_j) & \#scenario\,1, 2 \quad T(i, j) = 4 \end{cases} \tag{12}$$

, where both $\mu_{i-1}$ and $\mu_i$ are aligned with $s_j$ when $T(i, j) = 4$. This is the only difference between our recursion and the recursion of the standard global alignment algorithm.

Based on the proposed recursion, we developed two alignment algorithms designed for different problems: the full alignment algorithm (Sec. 5.1) allows insertions and deletions in the reference sequence, while the partial alignment algorithm (Sec. 5.2) does not allow any insertions or deletions in the partial reference sequence. The full alignment algorithm tries to align all the current signal segments to the whole reference sequence, while the partial alignment algorithm tries to align all the current signal segments to a consecutive sub-sequence of the 5-mer list from the beginning.

The output of the alignment algorithms is a matching path between the current signal segments $\boldsymbol{\mu}$ and the reference 5-mer list $\boldsymbol{s}$. We denote it by $\boldsymbol{\gamma} = (\boldsymbol{\mu}^{\boldsymbol{p}}, \boldsymbol{s}^{\boldsymbol{q}})$, where $\boldsymbol{\mu}^{\boldsymbol{p}} = (\mu_{p_1}, \mu_{p_2}, \cdots, \mu_{p_N})$ and $\boldsymbol{s}^{\boldsymbol{q}} = (s_{q_1}, s_{q_2}, \cdots, s_{q_N})$. The length of the alignment path is given by $|\boldsymbol{\gamma}| = N$. $\boldsymbol{p} = (p_1, p_2, \cdots, p_N)$ are the indexes of current signal segment $\boldsymbol{\mu}$ in the alignment path $\boldsymbol{\gamma}$, while $\boldsymbol{q} = (q_1, q_2, \cdots, q_N)$ are the indexes of the 5-mer list $\boldsymbol{s}$. Note that there are duplicated values in $\boldsymbol{p}$ and $\boldsymbol{q}$ to allow one 5-mer to match with multiple current signal segments. Fig. 4b provides an illustration of the alignment result.

To obtain the modification states $\boldsymbol{u}$, we will first process the optimal alignment path $\boldsymbol{\gamma}$ into $\hat{\boldsymbol{\gamma}}$ (Sec. 5.3), from which we can derive the modification states $\boldsymbol{u}$ (Sec. 6.1) .

#### 5.1 Full alignment algorithm

The full alignment algorithm is used when the read may possess differences (insertions or deletions) compared with the reference sequence, which is the general case. The algorithm is described in Algorithm 1.

**Algorithm 1:** Full alignment algorithm

**Input:** $\boldsymbol{\mu}$, $\boldsymbol{s}$, $\boldsymbol{T}^{kmer}$
**Output:** Alignment path $\boldsymbol{\gamma}$

```
// Add special symbol before μ and s
μ = {0.0, μ}, s = {"−", s}
// Get the length of μ and s
m = |μ|, n = |s|
// Initialize the score matrix M and traceback matrix T
Initialize a m × n matrix M, with all elements set to 0.0
Initialize a m × n matrix T, with all elements set to 0
// Initialize the first row of M and T
for i ∈ {1, 2, ··· , n − 1} do
    M(0, i) ← M(0, i − 1) + f(0.0, s_i)
    T(0, i) ← 2
end
// Intialize the first column of M and T
for i ∈ {1, 2, ··· , m − 1} do
    M(i, 0) ← M(i − 1, 0) + f(μ_i, "−")
    T(i, 0) ← 3
end
// Loop for all matrix
for i ∈ {1, 2, ··· , m − 1} do
    for j ∈ {1, 2, ··· , n − 1} do
        v_1 ← M(i − 1, j − 1) + f(μ_i, s_j)
        v_2 ← M(i, j − 1) + f(0.0, s_j)
        v_3 ← M(i − 1, j) + f(μ_i, "−")
        v_4 ← M(i − 1, j) + f(μ_i, s_j)
        M(i, j) ← max{v_1, v_2, v_3, v_4}
        T(i, j) ← argmax{v_1, v_2, v_3, v_4} # index of maximum
    end
end
// Traceback
Traceback from position (m − 1, n − 1) to position (0, 0) using T to generate the optimal
  alignment path γ.
```

#### 5.2 Partial alignment algorithm

The partial alignment algorithm is used when the read must agree with the reference sequence (no insertions or deletions). If the sequencing data is generated by *in vitro transcription* from a single template, all reads should agree with the reference sequence. However, the read may only match the reference sequence from the start to a middle point due to the break of the RNA molecule in the reverse transcription.

The partial alignment algorithm is a modification of the full alignment algorithm by considering this special case. Since the alignment may not be full, we traceback from the maximum value in the last row of the scoring matrix $M$. This means all current signal segments must be aligned, while only the first part of the 5-mer list needs to be aligned. Fig. 4c provides an illustration of the alignment result.

The partial alignment algorithm is described in Algorithm 2. We can see that scenarios 3 and 4 have been removed from the algorithm compared with the full alignment algorithm, i.e. $T(i,j) = 2 or 3$. In other words, there should be no insertions or deletions in the 5-mer list $\boldsymbol{s}$.

**Algorithm 2:** Partial alignment algorithm

**Input:** $\boldsymbol{\mu}$, $\boldsymbol{s}$, $\boldsymbol{T}^{kmer}$
**Output:** Alignment path $\boldsymbol{\gamma}$
```
// Add special symbol before μ and s
μ = {0.0, μ}, s = {"−", s}
// Get the length of μ and s
m = |μ|, n = |s|
// Initialize the score matrix M and traceback matrix T
Initialize a m × n matrix M, with all elements set to 0.0
Initialize a m × n matrix T, with all elements set to 0
// Initialize the first row of M and T
for i ∈ {1, 2, ··· , n − 1} do
    M(0, i) ← M(0, i − 1) + f(0.0, s_i)
    T(0, i) ← 2
end
// Intialize the first column of M and T
for i ∈ {1, 2, ··· , m − 1} do
    M(i, 0) ← M(i − 1, 0) + f(μ_i, "−")
    T(i, 0) ← 3
end
// Loop for all matrix
for i ∈ {1, 2, ··· , m − 1} do
    for j ∈ {1, 2, ··· , n − 1} do
        v_1 ← M(i − 1, j − 1) + f(μ_i, s_j)
        v_4 ← M(i − 1, j) + f(μ_i, s_j)
        M(i, j) ← max{v_1, v_4}
        if v_1 ≥ v_4 then
            T(i, j) ← 1;
        else
            T(i, j) ← 4;
        end
    end
end
// Traceback
Find the maximum value M(m − 1, k) in the last row of the scoring matrix M.
Traceback from position (m − 1, k) to position (0, 0) using T to generate the optimal
  alignment path γ.
```

##### 5.3 Post-processing of alignment path

In the alignment path $\boldsymbol{\gamma}$, one $k$-mer might be aligned to noise (scenario 4) or one current signal segment may be aligned with "-" (scenario 3). Firstly we remove such pairs from the alignment path. Then, we re-compute a weighted mean for each remaining $k$-mer, since one $k$-mer may be aligned to multiple current signal segments. Here we combine these segments and calculate the weighted mean over corresponding data points with hidden state 1, i.e. $z_i = 1$.

Let us assume there are $m$ segments aligned to the same $k$-mer $s_i$, with $n_j$ denotes the number of data points assigned to "curr" state ($z_i = 1$) in the $j$-th segment and $\mu_j$ denotes the mean of the $j$-th segment. The weighted mean, denoted by $\hat{\mu}_i$, corresponding to the $k$-mer $s_i$, is defined as follows:

$$\hat{\mu}_i = \sum_{j=1}^{m} \frac{n_j}{n_1 + n_2 + \cdots + n_m} \mu_j. \tag{13}$$

In the end, we obtain a one-to-one correspondence between each $k$-mer and its newly derived weighted mean, denoted by $\hat{\boldsymbol{\mu}} = (\hat{\mu}_1, \hat{\mu}_2, \cdots)$. Note that the length of $\hat{\boldsymbol{\mu}}$ and $\boldsymbol{s}$ is the same, i.e. $|\hat{\boldsymbol{\mu}}| = |\boldsymbol{s}|$. We denote the post-processed alignment path by $\hat{\boldsymbol{\gamma}} = (\hat{\boldsymbol{\mu}}, \boldsymbol{s})$.

### 6 Modification estimation

##### 6.1 Modification state estimation on single molecule level

We are interested in estimating the modification state for 5-mer after alignment. $\boldsymbol{T}^{kmer}$ has provided the mean and std for each 5-mer, as well as its modifications. Note that $\boldsymbol{T}^{kmer}$ may only contain modification parameters for only a subset of 5-mers. For those 5-mers without modification parameters, its mean and std are set to "NA". We could evaluate the probability density of a given 5-mer under the unmodified distribution or the modified distribution. By comparing the likelihoods, we could infer the modification state. Given the post-processed alignment path $\hat{\boldsymbol{\gamma}} = (\hat{\boldsymbol{\mu}}, \boldsymbol{s})$, we estimate the modification state of $j$th 5-mer $s_j$ by

$$u_j = \begin{cases} \text{"}un\text{"} \quad if \quad \mu^{un}_{s_j} = \text{"}NA\text{"} \; or \; \sigma^{un}_{s_j} = \text{"}NA\text{"} \\ \text{"}un\text{"} \quad if \quad N\big(\hat{\mu}_j \mid \mu^{un}_{s_j}, (\sigma^{un}_{s_j})^2\big) \geq N\big(\hat{\mu}_j \mid \mu^{mod}_{s_j}, (\sigma^{mod}_{s_j})^2\big) \\ \text{"}mod\text{"} \quad if \quad N\big(\hat{\mu}_j \mid \mu^{un}_{s_j}, (\sigma^{un}_{s_j})^2\big) < N\big(\hat{\mu}_j \mid \mu^{mod}_{s_j}, (\sigma^{mod}_{s_j})^2\big) \end{cases} \tag{14}$$

where $(\mu^{un}_{s_j}, \sigma^{un}_{s_j}) = T^{kmer}_{s_j,un}$ and $(\mu^{mod}_{s_j}, \sigma^{mod}_{s_j}) = T^{kmer}_{s_j,mod}$.

In the end, we get the modification state vector $\boldsymbol{u} = (u_1, u_2, \cdots, u_i, \cdots, u_{|\hat{\boldsymbol{\gamma}}|})$ for each 5-mer in the post-processed alignment path $\hat{\boldsymbol{\gamma}}$.

##### 6.2 Modification probability estimation on single molecule level

In addition to estimating the modification state, it is also crucial to determine the modification probability of each 5-mer on each read, which is essential for accurately benchmarking the performance at the single-molecule level.

Given the post-processed alignment path $\hat{\boldsymbol{\gamma}} = (\hat{\boldsymbol{\mu}}, \boldsymbol{s})$, we estimate the modification probability of $j$th 5-mer $s_j$ by

$$p_j = \begin{cases} \text{"}NA\text{"} \quad if \quad \mu^{un}_{s_j} = \text{"}NA\text{"} \; or \; \sigma^{un}_{s_j} = \text{"}NA\text{"} \\ \dfrac{p(\hat{\mu}_j | T^{kmer}_{s_j,mod})}{p(\hat{\mu}_j | T^{kmer}_{s_j,mod}) + p(\hat{\mu}_j | T^{kmer}_{s_j,un})} \quad if \quad \mu^{un}_{s_j} \neq \text{"}NA\text{"} \; and \; \sigma^{un}_{s_j} \neq \text{"}NA\text{"} \end{cases} \tag{15}$$

where $(\mu^{un}_{s_j}, \sigma^{un}_{s_j}) = T^{kmer}_{s_j,un}$ and $(\mu^{mod}_{s_j}, \sigma^{mod}_{s_j}) = T^{kmer}_{s_j,mod}$. Indeed, the $p_j$ is the posterior probability given the distribution of modified state:

$$p_j = p(T^{kmer}_{s_j,mod} | \hat{\mu}_j) = \frac{p(\hat{\mu}_j | T^{kmer}_{s_j,mod}) \cdot p(T^{kmer}_{s_j,mod})}{p(\hat{\mu}_j | T^{kmer}_{s_j,mod}) \cdot p(T^{kmer}_{s_j,mod}) + p(\hat{\mu}_j | T^{kmer}_{s_j,un}) \cdot p(T^{kmer}_{s_j,un})} \tag{16}$$

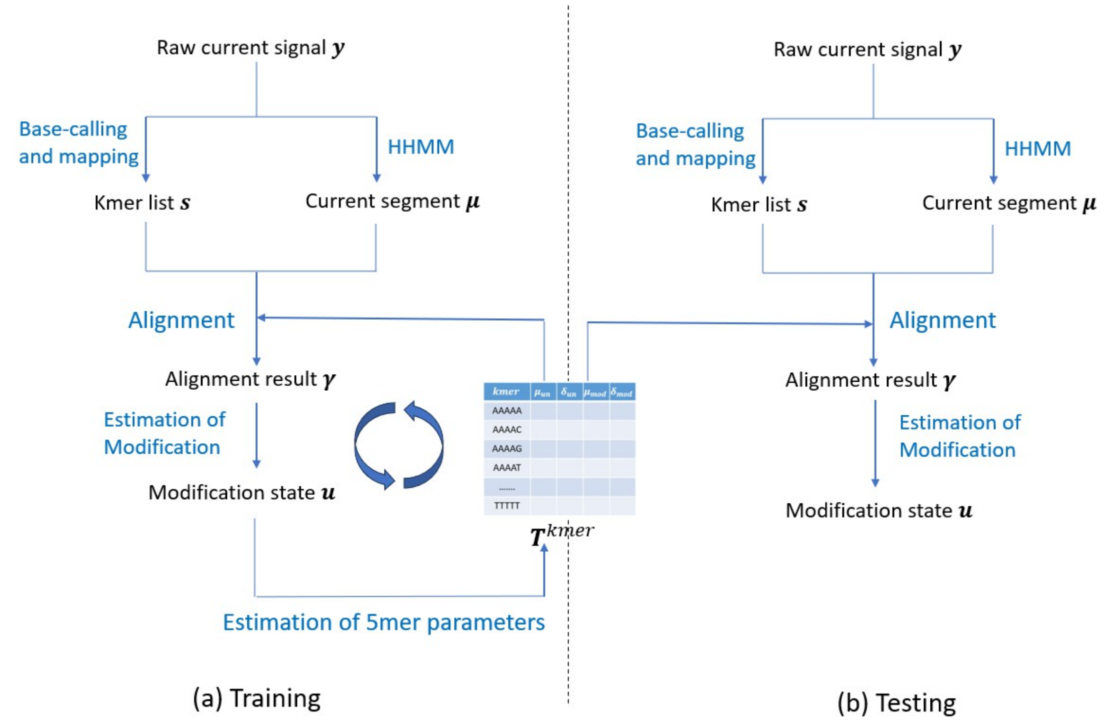

Figure 5: Estimation of 5-mer parameters. $\boldsymbol{T}^{kmer}$ is initialized using $\boldsymbol{T}^{ref}$. In training, our algorithm infers the parameters iteratively. After training, $\boldsymbol{T}^{ref}$ is fixed in testing.

where we assume that the prior probabilities are equal, $p(T^{kmer}_{s_j,mod}) = p(T^{kmer}_{s_j,un})$.

In the end, we get the modification probability vector $\boldsymbol{p} = (p_1, p_2, \cdots, p_i, \cdots, p_{|\hat{\boldsymbol{\gamma}}|})$ for each 5-mer in the post-processed alignment path $\hat{\boldsymbol{\gamma}}$.

##### 6.3 Modification rate estimation on site level

In real data analysis, we need to decide the modification rate of a genomic location, which quantifies the probability of a read mapped to this location to be in the modified state. For a genomic location, we denote the 5-mer from reads mapped to this location as $(s_1, s_2, \cdots, s_N)$. Note that all $s_i, i = 1, 2, \cdots, N$ are identical 5-mers on different reads. The modification state of these 5-mers on different reads are denoted by $(u_1, u_2, \cdots, u_N)$. The modification rate is given by

$$\eta = \frac{\sum_{i=1}^{N} u_i = \text{“}mod\text{”}}{N}$$

#### 7 Estimation of $k$-mer parameters table

This section provides the inference algorithms for 5-mer parameter table $\boldsymbol{T}^{kmer}$. This is necessary for training data, as we do not have accurate *prior* parameters for $\boldsymbol{T}^{kmer}$.

Given input data $\boldsymbol{Y} = (\boldsymbol{y_1}, \boldsymbol{y_2}, \cdots, \boldsymbol{y_i}, \cdots, \boldsymbol{y_N})$, we first perform basecalling and mapping to get $\boldsymbol{S} = (\boldsymbol{s_1}, \boldsymbol{s_2}, \cdots, \boldsymbol{s_i}, \cdots, \boldsymbol{s_N})$ run HHMM for all reads such we get $\boldsymbol{M} = (\boldsymbol{\mu_1}, \boldsymbol{\mu_2}, \cdots, \boldsymbol{\mu_i}, \cdots, \boldsymbol{\mu_N})$. We assume there are $n_i$ fragments for the $i$th read $\boldsymbol{y_i} = (\boldsymbol{y_i}^{(1)}, \boldsymbol{y_i}^{(2)}, \cdots, \boldsymbol{y_i}^{(n_i)})$. Similarly, we have $\boldsymbol{s_i} = (\boldsymbol{s_i}^{(1)}, \boldsymbol{s_i}^{(2)}, \cdots, \boldsymbol{s_i}^{(n_i)})$ and $\boldsymbol{\mu_i} = (\boldsymbol{\mu_i}^{(1)}, \boldsymbol{\mu_i}^{(2)}, \cdots, \boldsymbol{\mu_i}^{(n_i)})$.

The inference algorithm (Algorithm 3) then estimate $\boldsymbol{T}^{kmer}$ given $\boldsymbol{S}$, $\boldsymbol{M}$ and $\boldsymbol{T}^{ref}$ [4]. As illustrated in Fig. 5, the inference algorithm infers the parameters iteratively. With a given $\boldsymbol{T}^{kmer}$, we can use it to obtain the pairing between raw current signal segments and 5-mers, as well as modification states, through the alignment algorithm (Sec. 5). Given this pairing, we extract all current signal segments

[4] https://github.com/nanoporetech/kmer_models

for a specific 5-mer, which may correspond to different genomic locations and different reads. As there are two states for this 5-mer (unmodified and modified), we expect to observe two peaks in the density of the current signal segments. Therefore, we could fit a two-component Gaussian mixture model to the current signal segments to estimate relevant parameters of the given 5-mer in $\boldsymbol{T}^{kmer}$.

---

**Algorithm 3:** Inference algorithm of $k$-mer parameter table

---

**Input:** $\boldsymbol{S}$, $\boldsymbol{M}$, $\boldsymbol{T}^{ref}$
**Output:** $\boldsymbol{T}^{kmer}$

```
// Initialize T^kmer, ∀s ∈ {AAAAA, AAAAC, ⋯, TTTTT}, ∀u ∈ {"un", "mod"}
```

**for** $(\mu_{s,u}, \sigma_{s,u}) \in \boldsymbol{T}^{kmer}$ **do**
  $(\mu_{s,u}, \sigma_{s,u}) = \boldsymbol{T}^{ref}_{s,\text{"}un\text{"}}$
**end**

```
// iterate T times to update T^kmer
```

**for** $round = 1, 2 \cdots T$ **do**

```
  // align all fragment of all reads
```

  Perform full alignment algorithm (Alg. 1) for $\boldsymbol{\mu}_{\boldsymbol{i}}^{(k)}$ and $\boldsymbol{s}_{\boldsymbol{i}}^{(k)}$ to get the alignment path $\boldsymbol{\gamma}_{\boldsymbol{i}}^{(k)}$, where $i = 1 \cdots N$ and $k = 1 \cdots n_i$.
  Post-process (Sec. 5.3) the alignment path $\boldsymbol{\gamma}_{\boldsymbol{i}}^{(k)}$ into $\hat{\boldsymbol{\gamma}}_{\boldsymbol{i}}^{(k)} = [(\hat{\mu}_1^{(k)}, s_1^{(k)}), (\hat{\mu}_2^{(k)}, s_2^{(k)}), \cdots]$.
  Concatenate all $\hat{\boldsymbol{\gamma}}_{\boldsymbol{i}}^{(k)}$ and re-index into one vector $\hat{\boldsymbol{\Gamma}} = [(\hat{\mu}_1, s_1), (\hat{\mu}_2, s_2), \cdots]$.

```
  // parameter estimation
```

  **for** $s_0 \in \{AAAAA, AAAAC, \cdots, TTTTT\}$ **do**
    Collect all $\hat{\mu}_j$ in $\hat{\boldsymbol{\Gamma}}$ where $s_j = s_0$ into one vector $\boldsymbol{x} = \{\hat{\mu}_j,\ \forall s_j = s_0,\ \forall (\hat{\mu}_j, s_j) \in \hat{\boldsymbol{\Gamma}}\}$.
    Fit $\boldsymbol{x}$ using two component Gaussian Mixture Model with the first component mean fixed to $\mu_{s_0}$, where $(\mu_{s_0}, \sigma_{s_0}) = \boldsymbol{T}^{ref}_{s_0,\text{"}un\text{"}}$, and obtain the std of the first component $\sigma_1$, the mean and std of the second component $\mu_2$, $\sigma_2$.
    Update the parameters of 5-mer $s_0$ in $\boldsymbol{T}^{kmer}$: $\boldsymbol{T}^{kmer}_{s_0,\text{"}un\text{"}} \leftarrow (\mu_{s_0}, \sigma_1)$ and $\boldsymbol{T}^{kmer}_{s_0,\text{"}mod\text{"}} \leftarrow (\mu_2, \sigma_2)$
  **end**
  Update $\boldsymbol{T}^{kmer}$ manually based on several heuristic criteria.
**end**

---

In practice, the following heuristic criteria are used: (1) for a given 5-mer, perform GMM for each genomic location with the same 5-mer and only pool those sites with high modification rate (2) there must be two peaks in the density plot of a given 5-mer; (3) the mean of the two peaks are far apart; (4) the weight of second component (modified state) must be large, e.g. >0.1; (5) the weight of the first component (unmodified state) should be larger than the second component (modified state), as majority nucleotides are unmodified; (6) limit the size of std for both components in the GMM ($\sigma_1 < 5$, $\sigma_2 < 5$) to comply with the physical settings.

In the end, we will get an updated $\boldsymbol{T}^{kmer}$ from the training data, which contains the modification parameters for the strongest 5-mers. As illustrated in Fig. 2, we do have parameters for the modification state for certain $k$-mers such as "AAAAA", "AAAGC" etc.

### 8 Evaluation of segmentation and event alignment

The output of SegPore `eventalign` is the alignment path $\boldsymbol{\gamma} = (\boldsymbol{\mu}, \boldsymbol{s})$, which represents a one-to-one correspondence between the mean list $\boldsymbol{\mu} = (\mu_1, \mu_2, \ldots)$ and the $k$-mer list $\boldsymbol{s} = (s_1, s_2, \ldots)$. In addition, the corresponding standard deviation list $\boldsymbol{\sigma} = (\sigma_1, \sigma_2, \ldots)$ is also obtained along the alignment path. Fig. 6 is an example of the SegPore `eventalign` output, which illustrates one alignment path. The column `ref_kmer` corresponds to $\boldsymbol{s}$, the column `mean` corresponds to $\boldsymbol{\mu}$, and the column `stdv` corresponds to $\boldsymbol{\sigma}$.

Given the eventalign results, we use two metrics to evaluate the performance of the segmentation and event alignment : (1) the average std $\hat{\sigma}$, and (2) the average log-likelihood $\hat{L}$. Assuming there are $N$ reads in total (each read has one path), and the $n$th read has $K_n$ events ($|\boldsymbol{\gamma}_n| = K_n$). The average std $\hat{\sigma}$ is defined as

| read_idx | read_name | contig | pos | strand | ref_kmer | model_kmer | mean | stdv | start_idx | end_idx | event_len |
|---|---|---|---|---|---|---|---|---|---|---|---|
| 80741 | 003655f4-ff77-409d-a29e-20dc778d1c47 | A1 | 57 | + | GGTGT | GGTGT | 79.339 | 2.414 | 47296 | 47321 | 18 |
| 80741 | 003655f4-ff77-409d-a29e-20dc778d1c47 | A1 | 58 | + | GTGTC | GTGTC | 89.809 | 3.579 | 47276 | 47292 | 14 |
| 80741 | 003655f4-ff77-409d-a29e-20dc778d1c47 | A1 | 59 | + | TGTCT | TGTCT | 117.017 | 6.287 | 47251 | 47272 | 18 |
| 80741 | 003655f4-ff77-409d-a29e-20dc778d1c47 | A1 | 60 | + | GTCTT | GTCTT | 76.448 | 1.038 | 47148 | 47247 | 87 |
| 80741 | 003655f4-ff77-409d-a29e-20dc778d1c47 | A1 | 61 | + | TCTTA | TCTTA | 76.036 | 1.044 | 47117 | 47144 | 25 |
| 80741 | 003655f4-ff77-409d-a29e-20dc778d1c47 | A1 | 62 | + | CTTAG | CTTAG | 78.886 | 1.127 | 47092 | 47113 | 21 |
| 80741 | 003655f4-ff77-409d-a29e-20dc778d1c47 | A1 | 63 | + | TTAGT | TTAGT | 92.971 | 2.307 | 47007 | 47088 | 73 |
| 80741 | 003655f4-ff77-409d-a29e-20dc778d1c47 | A1 | 64 | + | TAGTG | TAGTG | 121.129 | 4.2 | 46882 | 46983 | 90 |
| 80741 | 003655f4-ff77-409d-a29e-20dc778d1c47 | A1 | 65 | + | AGTGT | AGTGT | 90.22 | 4.554 | 46822 | 46878 | 33 |
| 80741 | 003655f4-ff77-409d-a29e-20dc778d1c47 | A1 | 67 | + | TGTGC | TGTGC | 104.566 | 2.563 | 46798 | 46818 | 8 |
| 80741 | 003655f4-ff77-409d-a29e-20dc778d1c47 | A1 | 68 | + | GTGCT | GTGCT | 91.513 | 2.51 | 46778 | 46794 | 15 |
| 80741 | 003655f4-ff77-409d-a29e-20dc778d1c47 | A1 | 69 | + | TGCTT | TGCTT | 111.485 | 4.001 | 46754 | 46774 | 10 |

Figure 6: SegPore eventalign output example

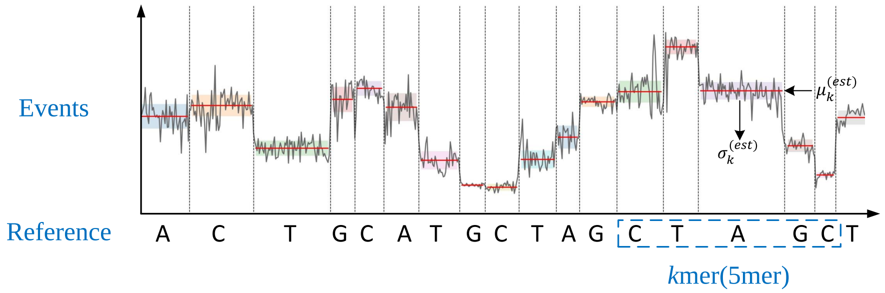

Figure 7: Illustration of segmentation and event alignment

$$\hat{\sigma} = \frac{1}{N} \sum_{n=1}^{N} \left\{ \frac{1}{K_n} \sum_{k=1}^{K_n} \sigma_k \right\}, \tag{17}$$

and the average log-likelihood $\hat{L}$ is defined as

$$\hat{L} = \frac{1}{N} \sum_{n=1}^{N} \left\{ \frac{1}{K_n} \sum_{k=1}^{K_n} \log \mathcal{N}\Big( \mu_k | \mu_{s_k}^{ref}, \sigma_{s_k}^{ref} \Big) \right\} \tag{18}$$

where $(\mu_{s_k}^{ref}, \sigma_{s_k}^{ref}) = T_{s_k}^{ref}$.

As shown in Fig. 7, the red line represents the event mean $\mu_k$ and the shaded area represents the std $\sigma_{s_k}^{est}$ for event $k$. A poorly segmented raw signal corresponding to an event will exhibit a large standard deviation. Therefore, a smaller $\hat{\sigma}$ indicates lower variations within the raw signal segment, signifying better performance.

If an event is aligned to the correct reference $k$-mer, the mean $\mu_k$ will be close to the reference $\mu_{s_k}^{ref}$ and the log-likelihood $\hat{L}$ will be large. So higher $\hat{L}$ means more similar results to ONT's estimates and better performances.

### GPU-accelerated Hierarchical Hidden Markov Model parameter inference

(Supplementary Note 2)

Guangzhao Cheng, Aki Vehtari, Lu Cheng

October, 2025

#### Contents

**1 Introduction** 1

**2 Enumeration of hidden states of outer HMM** 2

**3 General inference algorithm** 3

**4 Implementation details** 5

#### 1 Introduction

We have introduced the hierarchical hidden Markov model (HHMM) in **Supplementary Note 1**. The HHMM is a two-layer HMM, consisting of the outer HMM and the inner HMM. The outer HMM specifies the base blocks and transition blocks. The inner HMM specifies the base blocks. Linear models are used for the transition blocks. However, we have not provided the details for the parameter inference. This document aims to provide a comprehensive introduction to the inference algorithm.

As we know, the raw fast5 files are very large, which contain millions of reads. Each read has 20,000 measurements in the raw current signal. This poses a significant computational burden to the HHMM inference algorithm. Therefore, we used GPU to accelerate the inference algorithm.

We will first introduce the general inference algorithm, then describe the implementation details.

For simplicity, we assume the input is raw current signal of a single read with $N$ measurements $\boldsymbol{y} = (y_1, y_2, \cdots, y_N)$. The output are the hidden states of the outer HMM $\boldsymbol{g} = (g_1, g_2, \cdots, g_N)$ and the hidden states of the inner HMM $\boldsymbol{h} = (h_1, h_2, \cdots, h_N)$, where $g_i \in \{\text{“}B\text{”}, \text{“}T\text{”}\}$ and $h_i \in \{\text{“}curr\text{”}, \text{“}prev\text{”}, \text{“}next\text{”}, \text{“}noise\text{”}\}$.

Given a hidden state configuration of the outer HMM $\boldsymbol{g}$, it partitions the raw current signal $\boldsymbol{y}$ into alternating base blocks and transition blocks $\boldsymbol{c} = (c_1, c_2, \cdots, c_i, \cdots, c_{2K+1})$, where $c_i \in \{\text{“}B\text{”}, \text{“}T\text{”}\}$ denotes the label of $i$th block. Note that $c_i = \text{“}B\text{”}\, \forall i = 1, 3, \cdots, 2K+1$ and $c_i = \text{“}T\text{”}\, \forall i = 2, 4, \cdots, 2K$. Now we can denote $\boldsymbol{y} = (\boldsymbol{y^{(1)}}, \boldsymbol{y^{(2)}}, \cdots, \boldsymbol{y^{(2K+1)}})$.

The general idea of the inference algorithm is that we will calculate the likelihood for all possible configurations of the hidden states of the outer HMM $\boldsymbol{g}$ and choose the one with the largest likelihood as our final estimation. Due to the special constraints of $\boldsymbol{g}$, we can enumerate all possible configurations (Sec. 2). Here we denote the full configuration set of $\boldsymbol{g}$ by $\mathcal{G}$. The general inference algorithm is discussed in detail in Sec. 3.

The general inference algorithm (Sec. 3) provides the analytic forms for calculating the exact likelihood. However, a direct implementaion of the alogithm (Python and C++) requires huge amounts of computation resources and time to handle a typical fast5 file. Therefore, we implement a GPU version of the inference algorithm (Sec. 4), in which base blocks from different configurations are simultaneously inferred on different GPU cores.

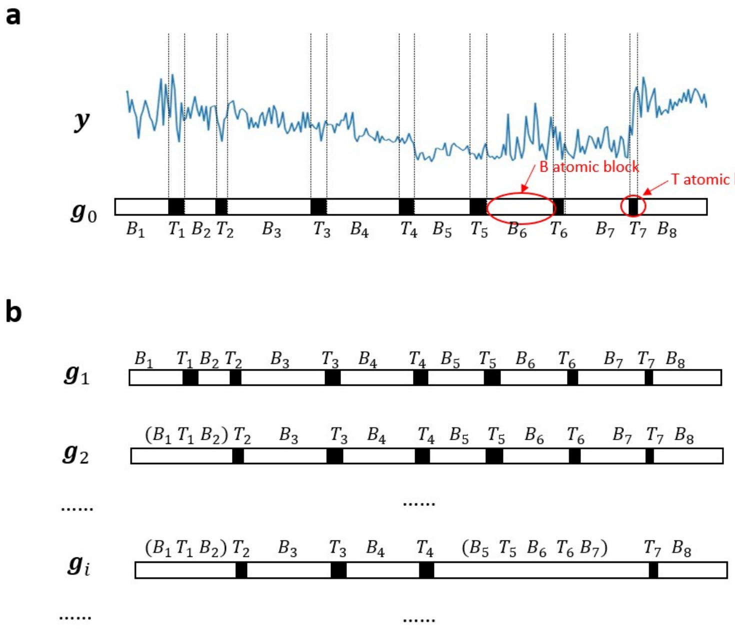

Figure 1: Illustration of enumeration of hidden states of outer HMM. (a) Raw signal and initialized to atomic segmentations. (b) Examples of the full configuration set $\mathcal{G} = (\boldsymbol{g}_1, \boldsymbol{g}_2, \cdots, \boldsymbol{g}_i, \cdots, \boldsymbol{g}_M)$.

### 2 Enumeration of hidden states of outer HMM

This section describes how to get the full configuration set $\mathcal{G} = (\boldsymbol{g}_1, \boldsymbol{g}_2, \cdots, \boldsymbol{g}_i, \cdots, \boldsymbol{g}_M)$ given the raw signal $\boldsymbol{y}$. Obviously, it is not possible to enumerate all hidden state $\boldsymbol{g}_i = (g_{i1}, g_{i2}, \cdots, g_{ij}, \cdots, g_{iN})$ (where $g_{ij} \in \{``B", ``T"\}$) because there are $2^N$ configurations and $N$ can be larger than $20,000$. However, there exist special properties of the raw signal $\boldsymbol{y}$: (1) the signal should be in alternating base and transition blocks (2) the hidden state $\boldsymbol{g}_i$ does not change within a block (3) the average length of base blocks is larger than that of the transition blocks (4) the variation of base blocks is smaller than that of transition blocks. Based on these data properties, we could partition the raw signal $\boldsymbol{y}$ into atomic level base and transition blocks (Sec. 4.1). Multiple consecutive atomic level base and transition blocks can be merged into a single larger base block. As shown in Fig. 1a, we partition the raw signal $\boldsymbol{y}$ into 15 atomic-level blocks $(B1, T1, B2, T2, \cdots, B7, T7, B8)$. We can merge $(B1, T1, B2)$ into a new base block or $(B5, T5, B6, T6, B7)$ into a new base block (Fig. 1b).

Given a pre-defined partition $\boldsymbol{g}_0$ of raw signal $y$, we can derive the $2K+1$ atomic blocks $(B_1, T_1, B_2, T2, \cdots, B_K, T_K, B_{K+1}))$. To facilitate the description of the enumeration algorithm, we define $seg(i,j) = (B_{i+1}, T_{i+1}, B_{i+2}, \cdots, T_{j-1}, B_j)$ to be the merged block of all blocks in $seg(i,j)$, where $0 \leq i < j \leq K+1$. A path is then a list of non-overlapping and consecutive blocks. For example, the path corresponding to $\boldsymbol{g}_i$ in Fig. 1b is given by $[seg(0,2), T_2, B_3, T_3, B_4, T_4, seg(4,7), T_7, B_8]$. To reduce the number of paths, we set the maximum number of base blocks in any merged block to $N_{max}$. The enumeration algorithm is given by Algorithm 1.

| | "B" | "T" |
|---|---|---|
| "B" | 0.99 | 0.01 |
| "T" | 0.10 | 0.90 |

Table 1: Transition parameters of the outer HMM ($T^{outer}$).

| | "curr" | "prev" | "next" | "noise" |
|---|---|---|---|---|
| "curr" | 0.925 | 0.025 | 0.025 | 0.025 |
| "prev" | 0.300 | 0.500 | 0.100 | 0.100 |
| "next" | 0.300 | 0.100 | 0.500 | 0.100 |
| "noise" | 0.300 | 0.100 | 0.100 | 0.500 |

Table 2: Transition parameters of the inner HMM ($T^{inner}$) .

**Algorithm 1:** Enumeration of hidden states of outer HMM

```
Input: g_0, N_max, K
Output: 𝒢
Derive (B_1, T_1, B_2, T2, ··· , B_K, T_K, B_{K+1})) from g_0
𝒢 ← { }
for path in recur_enum(0) do
    Convert path to g
    Append g to 𝒢
end
// Recursive function to enumerate all paths starting from position st
Function recur_enum(st):
    res = [ ]
    if st = K then
        return [B_{K+1}]
    end
    // "[" is inclusive and ")" is exclusive
    for i ∈ [st, K + 1) do
        if i − st > N_max then
            return res
        end
        tmp_seg = seg(st, i)
        all_paths_from_i = recur_enum(i)
        for path in all_paths_from_i do
            tmp_path = concatenate tmp_seg with path
            Append tmp_path to res
        end
    end
    return res;
```

### 3 General inference algorithm

Given all possible hidden state configurations of the outer HMM $\mathcal{G}$, we want to calculate the likelihood for all $\boldsymbol{g} \in \mathcal{G}$. In order to calculate the likelihood, we need to specify the parameters for the emission distributions.

Given $\boldsymbol{g}$, we can derive that $\boldsymbol{y}$ is divided into $2K + 1$ alternating base and transition blocks. We denote all blocks by $\boldsymbol{y} = (\boldsymbol{y}^{(1)}, \boldsymbol{y}^{(2)}, \cdots, \boldsymbol{y}^{(i)}, \cdots, \boldsymbol{y}^{(2K+1)})$, where $\boldsymbol{y}^{(i)}$ represent the $i$th block. The base blocks are given by $(\boldsymbol{y}^{(b_1)}, \boldsymbol{y}^{(b_2)}, \cdots, \boldsymbol{y}^{(b_i)}, \cdots, \boldsymbol{y}^{(b_{K+1})})$, where $b_i = 2i-1$ and $i \in \{1, 2, \cdots, K+1\}$. The transition blocks are given by $(\boldsymbol{y}^{(t_1)}, \boldsymbol{y}^{(t_2)}, \cdots, \boldsymbol{y}^{(t_i)}, \cdots, \boldsymbol{y}^{(t_K)})$, where $t_i = 2i$ and $i \in \{1, 2, \cdots, K\}$.

We pre-specify the transition probabilities $T^{outer} = \{T^{outer}_{BB}, T^{outer}_{BT}, T^{outer}_{TB}, T^{outer}_{TT}\}$ between the base state and the transition state in the outer HMM, as shown in Table 1. The pre-specification has two considerations: (1) the probability of a transition from B to T after 10 consecutive B states is 0.1; (2) the probability of a transition from T to B after 10 consecutive T states is larger than 0.9; (3) to reduce computation load in the parameter inference. We pre-specify the initial probability of the first hidden state to 1, since $\boldsymbol{g}$ always starts with a "B" state.

For inner HMM, we pre-specify the transition probabilities $T^{inner}$ shown in Table 2, which is based on the following consideration (1) large proportion of the data should come from the "curr" state, i.e. it is easy to transit from other states to "curr" state; (2) it is unlikely to transit from "prev" to "next", and vice versa; (3) its is unlikely to transit from other states to the "noise" state. We pre-specify the initial probability of the first hidden state to 0.25, which gives equal probabilities for all four states. We pre-specify the parameters of the "noise" state ($Unif(lb, ub)$) to $lb = 50$ and $ub = 130$, which are estimated from the current signals across different datasets to represent a reasonable range.

The remaining parameters, mainly associated with emission distributions, need to be estimated. For base blocks, we have inner HMM parameters $\boldsymbol{\phi} = (\phi^{(b_1)}, \phi^{(b_2)}, \cdots, \phi^{(b_{K+1})})$ and $\boldsymbol{\sigma} = (\sigma^{(b_1)}, \sigma^{(b_2)}, \cdots, \sigma^{(b_{K+1})})$. For transition blocks, we have linear coefficients $\boldsymbol{\beta}_0 = (\beta_0^{(t_1)}, \beta_0^{(t_2)}, \cdots, \beta_0^{(t_K)})$, $\boldsymbol{\beta}_1 = (\beta_1^{(t_1)}, \beta_1^{(t_2)}, \cdots, \beta_1^{(t_K)})$ and noise std $\boldsymbol{\sigma_\epsilon} = (\sigma_\epsilon^{(t_1)}, \sigma_\epsilon^{(t_2)}, \cdots, \sigma_\epsilon^{(t_K)})$.

The general idea of the inference algorithm is to calculate the joint likelihood of the outer HMM given for any $g$. However, it is not possible to calcualte the emission probability for each measurement as there exists dependencies with a base or transition block. Here we first calculate the joint likelihood for a base block (inner HMM) or a transition block (linear model), then average the joint likelihood to each measurement to use it as substitue for the emission probability of each measurement in the outer HMM. Here we use $\boldsymbol{\omega} = (\omega_1, \omega_2, \cdots, \omega_N)$ to denote the approximated emission probability of each measurement. Note that $\omega_i$ is derived either based on the inner HMM or the linear model, and $|\boldsymbol{\omega}| = |\boldsymbol{y}| = |\boldsymbol{g}| = |\boldsymbol{h}|$.

The general inference algorithm is given by Algorithm 2.

**Algorithm 2:** General inference algorithm for HHMM

**Input:** $\boldsymbol{y}$, $\mathcal{G}$, $T^{outer}$, $\boldsymbol{\pi}^{outer}$, $T^{inner}$, $\boldsymbol{\pi}^{inner}$,

**Output:** best path $\tilde{\boldsymbol{g}}$ of outer HMM, best path $\tilde{\boldsymbol{h}}$ of inner HMM, $\boldsymbol{\phi}$, $\boldsymbol{\sigma}$, $\boldsymbol{\beta}_0$, $\boldsymbol{\beta}_1$, $\boldsymbol{\sigma_\epsilon}$

`// Initialization`

set all elements of $\tilde{\boldsymbol{\omega}}$, $\tilde{\boldsymbol{g}}$ and $\tilde{\boldsymbol{h}}$ to 0, and $|\tilde{\boldsymbol{g}}| = |\tilde{\boldsymbol{h}}| = |\boldsymbol{y}|$

$\mathcal{H} \leftarrow \{\}$

$\Gamma \leftarrow \{\}$

`// Calculate likelihood for each` $\boldsymbol{g}$

**for** $\boldsymbol{g} \in \mathcal{G}$ **do**

- $\boldsymbol{\omega} \leftarrow \tilde{\boldsymbol{\omega}}$
- $\boldsymbol{h} \leftarrow \tilde{\boldsymbol{h}}$
- Divide $\boldsymbol{y}$ into base and transition blocks $(\boldsymbol{y}^{(1)}, \boldsymbol{y}^{(2)}, \cdots, \boldsymbol{y}^{(2K+1)})$ based on $\boldsymbol{g}$
- `// transition state parameter estimation`
- Estimate $\hat{\boldsymbol{\omega}}^{(t_i)}$, $\beta_0^{(t_i)}$, $\beta_1^{(t_i)}$,$\sigma_\epsilon{}^{(t_i)}$ using linear model (Alg. 3), where $t_i = 2i$ and $i \in \{1, 2, \cdots, K\}$
- Update relevant part of $\boldsymbol{\omega}$ using $\hat{\boldsymbol{\omega}}^{(t_i)}$.
- `// base state parameter estimation`
- Initialize $\phi^{(b_i)}$ and $\sigma^{(b_i)}$ use the empirical mean and std of $\boldsymbol{y}^{(b_i)}$, where $b_i = 2i - 1$ and $i \in \{1, 2, \cdots, K+1\}$
- $\boldsymbol{\phi}^{new} \leftarrow \boldsymbol{\phi}$
- $\boldsymbol{\sigma}^{new} \leftarrow \boldsymbol{\sigma}$
- **for** *round* $= 1, 2, \cdots, R$ **do**
  - **for** $i \in (1, 2, \cdots, K+1)$ **do**
    - $b_i \leftarrow 2i - 1$
    - `// inner HMM parameter estimation`
    - Estimate $\hat{\boldsymbol{\omega}}^{(b_i)}$, $\hat{\boldsymbol{h}}^{(b_i)}$, $\hat{\phi}^{(b_i)}$, $\hat{\sigma}^{(b_i)}$ using Forward-backward algorithm (Alg. 4) given $\boldsymbol{y}^{(b_i)}$, $\phi^{(b_i)}$, $\sigma^{(b_i)}$, $\phi^{(b_{i-1})}$, $\sigma^{(b_{i-1})}$, $\phi^{(b_{i+1})}$, $\sigma^{(b_{i+1})}$, $T^{inner}$, $\boldsymbol{\pi}^{inner}$
    - Update corresponding elements of $\boldsymbol{\phi}^{new}$ and $\boldsymbol{\sigma}^{new}$ using $\hat{\phi}^{(b_i)}$, $\hat{\sigma}^{(b_i)}$
    - Update relevant part of $\boldsymbol{h}$ and $\boldsymbol{\omega}$ using $\hat{\boldsymbol{h}}^{(b_i)}$ and $\hat{\boldsymbol{\omega}}^{(b_i)}$
  - **end**
  - $\boldsymbol{\phi} \leftarrow \boldsymbol{\phi}^{new}$
  - $\boldsymbol{\sigma} \leftarrow \boldsymbol{\sigma}^{new}$
- **end**
- Calculate the joint likelihood of the outer HMM using $\boldsymbol{g}$, $\boldsymbol{\omega}$, $T^{outer}$, $\boldsymbol{\pi}^{outer}$

$$\gamma = \pi_{g_1}^{outer} \prod_{i=2}^{N} T_{g_{i-1}g_i}^{outer} \prod_{j=1}^{N} \boldsymbol{\omega}_j$$

- Append $\boldsymbol{h}$ and $\gamma$ to $\mathcal{H}$ and $\Gamma$, respectively.

**end**

Choose the largest likelihood from $\Gamma$ and retrieve the corresponding $\boldsymbol{g}$ and $\boldsymbol{h}$ from $\mathcal{G}$ and $\mathcal{H}$, respectively.

$\tilde{\boldsymbol{g}} \leftarrow \boldsymbol{g}$

$\tilde{\boldsymbol{h}} \leftarrow \boldsymbol{h}$

### 4 Implementation details

#### 4.1 Atomic block initialization

As mentioned in Sec. 2, we need to partition the raw signal $\boldsymbol{y}$ into $2K+1$ atomic blocks $(B_1, T_1, B_2, T2, \cdots, B_K, T_K, B_{K+1}))$. If we can identify all atomic transition blocks $(T_1, T_2, \cdots, T_K)$, then the atomic base blocks are automatically determined.

We assume atomic transition blocks $(T_1, T_2, \cdots, T_K)$ have higher variations than atomic base blocks. If we fit a line to a transition block, the absolute value of the slope of the line should be large. Based on this idea, we can obtain the atomic transition blocks as follows:

1. For each measurement $i$, fit another line to $(y_{i-1}, y_i, y_{i+1})$ and obtain the slope $\beta_1$, fit a line to $(y_{i-2}, y_{i-1}, y_i, y_{i+1}, y_{i+2})$ and obtain the slope $\beta_2$, use $\beta_i = |0.5 * (\beta_1 + \beta_2)|$ as the absolute slope for $y_i$.

2. Smooth the obtain slopes $\boldsymbol{\beta} = (\beta_1, \beta_2, \cdots, \beta_N)$ by taking the mean of a sliding window of size. Now we get the smoothed $\bar{\boldsymbol{\beta}}$.

3. Find peaks from $\bar{\boldsymbol{\beta}}$ using the `find_peaks()` function in the Python `Scipy.signal` package. Here set the minimum distance of detected peaks to 10. Now we get a list of peak positions $\boldsymbol{\theta} = (\theta_1, \theta_2, \cdots, \theta_K)$.

4. Expand $\theta_k$ to the atomic transition block $T_k$. We fit a 3-segment spline to $(y_{\theta_k-8}, \cdots, y_{\theta_k-l}, \cdots, y_{\theta_k}, \cdots, y_{\theta_k+r}, \cdots, y_{\theta_k+8})$, where $0 < l, r < 8$. The slopes of the first segment $(y_{\theta_k-8}, \cdots, y_{\theta_k-l-1})$ and the third segment $(y_{\theta_k+r+1}, \cdots, y_{\theta_k+8})$ are set to 0, while a standard linear model is fit to the second segment $(y_{\theta_k-l}, \cdots, y_{\theta_k}, \cdots, y_{\theta_k+r})$. In the fitting, the slope of the second segment should be larger than 5.0. After the fitting of the spline, we take the residuals and calculate the joint likelihood assuming each residual following a Gaussian distribution $N(0, 4)$. By enumeration all $l, r$, we pick the $\hat{l}, \hat{r}$ with the largest joint likelihood. Therefore, the final $k$th transition block $T_k$ corresponding to the interval $[\theta_k - l, \theta_k + r]$.

#### 4.2 Emission probabilities of linear model

For the $i$th transition block, the measurements is given by $\boldsymbol{y}^{(t_i)} = (y_1^{(t_i)}, y_2^{(t_i)}, \cdots, y_{N_{t_i}}^{(t_i)})$, where $t_i = 2i$ and $i \in 1, 2, \cdots, K$. We denote the corresponding index of $\boldsymbol{y}^{(t_i)}$ by $\boldsymbol{x}^{(t_i)} = (1, 2, \cdots, N_{t_i})$. Here we want to get the emission probabilities for each measurement $\hat{\boldsymbol{\omega}}^{(t_i)}$, which can be used by the outer HMM. Detail steps are provided in the following algorithm.

**Algorithm 3:** Emission probabilities of linear model

**Input:** $\boldsymbol{y}^{(t_i)}$, $\boldsymbol{x}^{(t_i)}$
**Output:** $\hat{\boldsymbol{\omega}}^{(t_i)}$, $\beta_0^{(t_i)}$, $\beta_1^{(t_i)}$,$\sigma_\epsilon^{(t_i)}$
initialize $\hat{\boldsymbol{\omega}}^{(t_i)}$ to a $1 \times N_{t_i}$ all zero vector.
Fit a linear model of $\boldsymbol{y}^{(t_i)}$ versus $\boldsymbol{x}^{(t_i)}$, obtain the maximum likelihood estimates $\beta_0^{(t_i)}$, $\beta_1^{(t_i)}$ and $\sigma_\epsilon^{(t_i)}$
Calculate the maximum log joint likelihood $\log p\big(\boldsymbol{y}^{(t_i)} | \boldsymbol{x}^{(t_i)}, \beta_0^{(t_i)}, \beta_1^{(t_i)}, \sigma_\epsilon^{(t_i)}\big) = \log N\big(\boldsymbol{y}^{(t_i)} - \beta_1^{(t_i)}\boldsymbol{x}^{(t_i)} - \beta_0^{(t_i)} \mid 0, (\sigma_\epsilon^{(t_i)})^2\big)$
Set each element of $\hat{\boldsymbol{\omega}}^{(t_i)}$ to $\exp\left(\frac{1}{N_{t_i}} \log p\big(\boldsymbol{y}^{(t_i)} | \boldsymbol{x}^{(t_i)}, \beta_0^{(t_i)}, \beta_1^{(t_i)}, \sigma_\epsilon^{(t_i)}\big)\right)$

#### 4.3 Emission probabilities of inner HMM

This section describes how to obtain the approximated emission probabilities of the outer HMM for a the $i$th base block $\boldsymbol{y}^{(b_i)} = (y_1^{(b_i)}, y_2^{(b_i)}, \cdots, y_{N_{b_i}}^{(b_i)})$, which is modeled using an inner HMM. Here $b_i = 2i-1$ and $i \in \{1, 2, \cdots, K\}$.

For the inner HMM, we list all the parameters in Table 3.

We want to obtain the approximated emission probabilities $\hat{\boldsymbol{\omega}}^{(b_i)}$, hidden states $\hat{\boldsymbol{h}}^{(b_i)}$, estimated parameters $\hat{\boldsymbol{\phi}}^{(b_i)}$ and $\hat{\boldsymbol{\sigma}}^{(b_i)}$ for the $i$th base block $\boldsymbol{y}^{(b_i)}$. To achieve this, we follow the Forward-backward algorithm in Bishop (2006) (Chapter 13.2 [BN06]). The general idea of the Forward-backward algorithm is to use a $Q$-function to approximate the marginal likelihood, then use an expectation-maximization (EM) algorithm for the parameter estimation.

We introduce the following notations for the Forward-backward algorithm. We use $\boldsymbol{Z} = (\boldsymbol{z}_1, \boldsymbol{z}_2, \cdots, \boldsymbol{z}_{N_{b_i}})$ to denote the hidden states, where $\boldsymbol{Z}$ is a $N_{b_i} \times 4$ matrix, $\boldsymbol{z}_j = (z_{j1}, z_{j2}, z_{j3}, z_{j4})$ and $z_{jk} \in \{0, 1\}$. If $z_{jk}$ is 1, then it means $h_j$ corresponds to the $k$th hidden state. We use $\boldsymbol{\alpha}$ ($N_{b_i} \times 4$) to denote the forward probabilities and $\boldsymbol{\beta}$ ($N_{b_i} \times 4$) to denote the backward probabilities. $\boldsymbol{\gamma}$ ($N_{b_i} \times 4$) denotes the posterior distribution calculated from $\boldsymbol{\alpha}$ and $\boldsymbol{\beta}$. $\boldsymbol{\xi}$ ($(N_{b_i} - 1) \times 4 \times 4$) denotes the joint distribution of two successive hidden states. We use $\boldsymbol{\theta} = (\phi^{(b_i)}, \sigma^{(b_i)})$ to denote all parameters to be estimated.

| Parameter | Fixed parameter | Description |
|---|---|---|
| $T^{inner}$ | Yes | Transition probabilities between hidden states (“curr”, “prev”, “next”, “noise”), specified in table 2 |
| $\pi_k^{inner}$ | Yes | probability of the first hidden state, $\pi_k^{inner} = 0.25$, where $k \in \{1, 2, 3, 4\}$ |
| $lb$ | Yes | lower bound of uniform distribution, $lb = 50$ |
| $ub$ | Yes | upper bound of uniform distribution, $ub = 130$ |
| $\phi^{(b_i)}$ | No | Gaussian mean of $i$th base block |
| $\sigma^{(b_i)}$ | No | Gaussian std of $i$th base block |
| $\phi^{(b_{i-1})}$ | No | Gaussian mean of $i-1$th base block |
| $\sigma^{(b_{i-1})}$ | No | Gaussian std of $i-1$th base block |
| $\phi^{(b_{i+1})}$ | No | Gaussian mean of $i+1$th base block |
| $\sigma^{(b_{i+1})}$ | No | Gaussian std of $i+1$th base block |

Table 3: Parameters of the inner HMM for the $b_i$th block.

$$
\begin{aligned}
Q(\boldsymbol{\theta}, \boldsymbol{\theta}^{old}) &= \sum_{\boldsymbol{Z}} p(\boldsymbol{Z}|\boldsymbol{y}^{(b_i)}, \boldsymbol{\theta}^{old}) \log p(\boldsymbol{y}^{(b_i)}, \boldsymbol{Z}|\boldsymbol{\theta}) \\
&= \sum_{k=1}^{4} \gamma_{1k} \log \pi_k^{inner} + \sum_{n=2}^{N_{b_i}} \sum_{j=1}^{4} \sum_{k=1}^{4} \xi_{n-1,j,k} \log T_{jk}^{inner} \\
&+ \sum_{n=1}^{N_{b_i}} \Big\{ \gamma_{n1} \log p(y_n^{(b_i)}|\phi^{(b_i)}, \sigma^{(b_i)}) + \gamma_{n2} \log p(y_n^{(b_i)}|\phi^{(b_{i-1})}, \sigma^{(b_{i-1})}) \\
&\qquad + \gamma_{n3} \log p(y_n^{(b_i)}|\phi^{(b_{i+1})}, \sigma^{(b_{i+1})}) + \gamma_{n4} \log p(y_n^{(b_i)}|lb, ub) \Big\}
\end{aligned} \tag{1}
$$

Here we only need to update $\boldsymbol{\alpha}$, $\boldsymbol{\beta}$, $\boldsymbol{\gamma}$, $\boldsymbol{\xi}$, $\phi^{(b_i)}$ and $\sigma^{(b_i)}$ in the inference process of inner HMM for $i$th base block. Our inference algorithm is a modification of the standard Forward-backward algorithm based on the defined $Q$-function, as described in Alg. 4.

$$
\phi^{(b_i)} = \frac{\sum_{n=1}^{N_{b_i}} \gamma_{n1} y_n^{(b_i)}}{\sum_{n=1}^{N_{b_i}} \gamma_{n1}} \tag{2}
$$

$$
\sigma^{(b_i)} = \frac{\sum_{n=1}^{N_{b_i}} \gamma_{n1} (y_n^{(b_i)} - \phi^{(b_i)})^2}{\sum_{n=1}^{N_{b_i}} \gamma_{n1}} \tag{3}
$$

**Algorithm 4:** Inference algorithm of inner HMM

**Input:** $\boldsymbol{y}^{(b_i)}$, $\phi^{(b_i)}$, $\sigma^{(b_i)}$, $\phi^{(b_{i-1})}$, $\sigma^{(b_{i-1})}$, $\phi^{(b_{i+1})}$, $\sigma^{(b_{i+1})}$, $T^{inner}$, $\boldsymbol{\pi}^{inner}$, $lb$, $ub$
**Output:** $\hat{\boldsymbol{\omega}}^{(b_i)}$, $\hat{\boldsymbol{h}}^{(b_i)}$, $\hat{\phi}^{(b_i)}$, $\hat{\sigma}^{(b_i)}$
Initialize $\hat{\boldsymbol{h}}^{(b_i)}$ and $\hat{\boldsymbol{\omega}}^{(b_i)}$ to the $1 \times N_{b_i}$ all zero vector.
Initialize $\boldsymbol{\alpha}$, $\boldsymbol{\beta}$, $\boldsymbol{\gamma}$, $\boldsymbol{\xi}$
`// EM algorithm`
**for** $round = 1, 2, \cdots, R$ **do**
  E-step: update $\boldsymbol{\alpha}$, $\boldsymbol{\beta}$, $\boldsymbol{\gamma}$ and $\boldsymbol{\xi}$ given $\boldsymbol{y}^{(b_i)}$, $\phi^{(b_i)}$, $\sigma^{(b_i)}$, $\phi^{(b_{i-1})}$, $\sigma^{(b_{i-1})}$, $\phi^{(b_{i+1})}$, $\sigma^{(b_{i+1})}$, $T^{inner}$, $\boldsymbol{\pi}^{inner}$, $lb$, $ub$
  M-step: calculate $\hat{\phi}^{(b_i)}$ and $\hat{\sigma}^{(b_i)}$ by Eq. 2 and Eq. 3
  $\phi^{(b_i)} \leftarrow \hat{\phi}^{(b_i)}$
  $\sigma^{(b_i)} \leftarrow \hat{\sigma}^{(b_i)}$
  Calculate the $Q$-function (log marginal likelihood) $\hat{q}$ given all estimated parameters
**end**
Derive $\hat{\boldsymbol{h}}^{(b_i)}$ from $\boldsymbol{\gamma}$ by taking the state with highest probability for each data point, i.e. $h_n^{(b_i)} = argmax([\gamma_{n1}, \gamma_{n2}, \gamma_{n3}, \gamma_{n4}])$
Set each element of $\hat{\boldsymbol{\omega}}^{(b_i)}$ to $\exp(\hat{q}/N_{b_i})$.

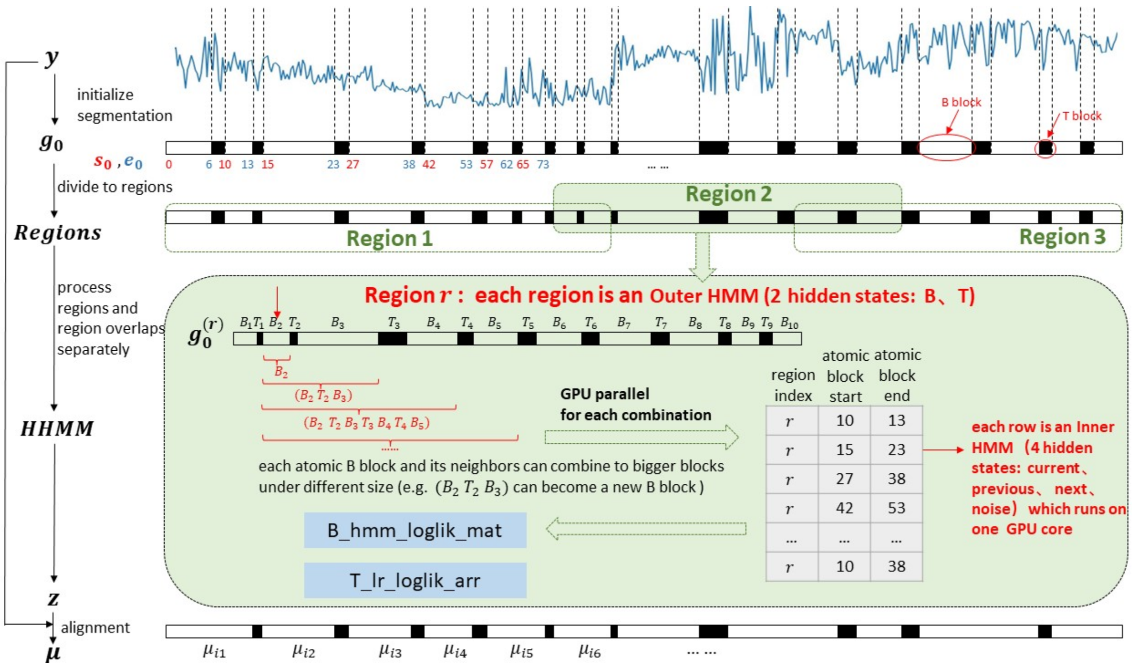

Figure 2: Illustration of GPU-accelerated parameter inference.

##### 4.4 GPU-accelerated parameter inference

Given the general inference algorithm (Sec. 3) and algorithms for calculating the emission probabilities for the inner HMM (Alg. 4) and linear model (Alg. 3), we have the full algorithm for the parameter inference. The actual inference, however, can be really slow since the input fast5 data are huge. Here we use heuristics and GPU parallelization to accelerate the parameter inference.

We divide the raw current signal of each read $\boldsymbol{y}$ into overlapping regions, which start at an atomic base block and also end with at an atomic base block. As shown in Fig. 2, we process the $r$th region $\boldsymbol{y}^{(r)}$ independently to get the segmentation results $\boldsymbol{z}^{(r)}$. This process is based on the following heuristics: the segmentation result of one region will not affect another region when they are far away from each other. The inference is performed in two rounds. In the first round, we obtain the segmentation results for non-overlapping regions. The center part of the segmentation is not affect by the flanks parts are affected may be affect by neighbouring regions. In the second round, we merge relevant parts of neighbouring regions near the touch point as a new region based on segmentation results of the first round. After that we perform the inference for the new regions to get the final segmentations for the whole read.

We use GPU to parallelize the inference algorithm of inner HMM (Alg. 4). For each region, we first enumerate all possible paths and calculate the likelihood for each candidate path (Sec. 2). As shown in Fig. 2, we need to perform inner HMM parameter inference for different paths of different regions of one read. We consider to parallel the computation of different merged blocks $seg(i, j)$ in different path of different regions. We first collect all merged blocks and sort them by size and only keep unique blocks, which can be identified by the combinatorial key of region index, start atomic block index, end atomic block index. Then we run the inner HMM inference (Alg. 4) of each merged block on a GPU core. Note that all GPU cores perform the same computation step of Alg. 4 simultaneously, but for different merged blocks.

#### References

[BN06] Christopher M Bishop and Nasser M Nasrabadi. *Pattern recognition and machine learning*, volume 4. Springer, 2006.
